# K5 polysaccharides inhibit SARS-CoV-2 infection by preventing spike-proteolytic priming

**DOI:** 10.1101/2025.04.23.650164

**Authors:** Maria Milanesi, Chiara Urbinati, Liv Zimmermann, Pasqua Oreste, Alberto Zani, Arnaldo Caruso, Francesca Caccuri, Vibor Laketa, Petr Chlanda, Rebecca C. Wade, Marco Rusnati, Giulia Paiardi

**Affiliations:** Macromolecular Interaction Analysis Unit, Section of Experimental Oncology and Immunology, Department of Molecular and Translational Medicine, 25123 Brescia, Italy; Schaller Research Group, Department of Infectious Diseases-Virology, Heidelberg University, 69120, Heidelberg, Germany; BioQuant, Heidelberg University, 69120, Heidelberg, Germany; Glycores 2000 S.r.l., 20115, Milan, Italy; Section of Microbiology, Department of Molecular and Translational Medicine, University of Brescia, 25123, Brescia, Italy; Department of Infectious Diseases, Virology, Medical Faculty Heidelberg, Heidelberg University, 69120 Heidelberg, Germany; German Center for Infection Research (DZIF), Partner Site Heidelberg, 69120 Heidelberg, Germany; Molecular and Cellular Modeling Group, Heidelberg Institute for Theoretical Studies (HITS), 69118 Heidelberg, Germany; Zentrum für Molekulare Biologie der Universität Heidelberg (ZMBH), DKFZ-ZMBH Alliance, 69120 Heidelberg, Germany; Interdisciplinary Center for Scientific Computing (IWR), Heidelberg University, 69120 Heidelberg, Germany; Consorzio Interuniversitario Biotecnologie (CIB), Unit of Brescia, 25123 Brescia, Italy

**Keywords:** antiviral agents, polysaccharides, K5, SARS-CoV-2, spike

## Abstract

SARS-CoV-2 spike glycoprotein is a promising drug target due to its crucial role in viral infection. Heparin, a long linear polysaccharide that inhibits SARS-CoV-2 infection by acting on spike, has limited antiviral applications due to its anticoagulant effect. *E. coli* K5 polysaccharides share the same structure as the heparin precursor and can be chemically modified to devoid anticoagulant activity. Here, biochemical assays and computer simulations reveal that K5 with a high degree of sulfation at O-(K5OSH) or N- and O-positions (K5NOSH) bind spike with higher affinity than heparin, preventing its binding to ACE2 and cleavage by furin. This mechanism is supported by a cell syncytia assay showing that K5OSH and K5NOSH inhibit viral infection by blocking membrane fusion. Infection assays for SARS-CoV-2 Wuhan-Hu-1 and Omicron BA.1 variants corroborate their antiviral activity. These results support the therapeutic potential of K5OSH and K5NOSH against SARS-CoV-2, with K5OSH displaying the more promising activity profile.

## Introduction

Since 2020, the COVID-19 pandemic caused by the severe acute respiratory syndrome coronavirus 2 (SARS-CoV-2) has heavily affected the lives of people all around the world. Despite the development of efficacious vaccines, surveillance systems, and personal protective equipment reducing the spread of infection, SARS-CoV-2 continues to result in serious symptoms, and it undergoes constant and rapid evolution of novel variants. Therefore, the development of broad-spectrum antiviral agents against SARS-CoV-2 is of high importance ^1^.

SARS-CoV-2 single-stranded positive-sense RNA virions are spherical with a diameter of 91±11 nm and composed of four main structural proteins: spike glycoprotein (spike), envelope (E), membrane (M), and nucleocapsid (N) protein ^2^. The spike is a homotrimeric class I fusion glycoprotein composed of S1 and S2 subunits that are respectively, responsible for the attachment of the virion to the host cell receptor via its receptor-binding domain (RBD) and the fusion with the host-cell membrane ^2, 3^. The RBD adopts an open-active or closed-inactive conformation. The RBD in the open-active conformation exposes the receptor binding motif (RBm), mediating the interaction with the host angiotensin-converting enzyme 2 (ACE2) receptor. Conversely, in the closed-inactive conformation, the RBm is shielded, hindering the binding to ACE2 ^4–7^. SARS-CoV-2 mediated membrane fusion requires sequential cleavage of S, which harbors a polybasic sequence that is cleaved at the S1/S2 site by furin during trafficking in a producer cell ^3, 8^. Once the spike has bound to the ACE2 receptor, it undergoes a second cleavage at the S2’ site by transmembrane protease serine 2 (TMPRSS2) at the plasma membrane or by cathepsins in the late endosomes. Upon S2’ cleavage, the S2 fusion peptide is liberated, and fusion of the viral and host membranes takes place ^3, 8–13^.

SARS-CoV-2 can infect a broad range of tissues and, depending on the available proteases, it either fuses with the plasma membrane or with the endosomal membrane. Endosomal SARS-CoV-2 entry is supported by heparan sulfate (HS) proteoglycans (HSPGs), co-receptors expressed on the host cell surface ^14–16^. HSPG consists of a core protein that carries multiple polysulfated glycosaminoglycans (GAG) chains. SARS-CoV-2 binding to host cell HSPGs is mainly driven by electrostatic forces involving the sulfate and carboxyl groups of HS and the positively charged amino acid residues in the spike protein basic domains. HS binds along a partially grooved basic channel on the spike spanning from the RBD in one subunit to the S1/S2 basic domain of the adjacent subunit ^17^. HS binding promotes the interaction of the spike with the host cell receptor ACE2, stabilizing the ternary complex and triggering structural rearrangement of the spike S2’ cleavage site ^17^. SARS-CoV-2 variants of concern (VOCs), in particular Omicron BA.1, have an increased net charge in spike, leading to enhanced dependence on HS for viral attachment and infection ^17–20^. Treatment of cells with HS structural analogs, such as unfractionated heparin or low molecular weight heparin and more generally, polysulfated GAGs, substantially reduces SARS-CoV-2 infection ^21–23^. This is explained by heparin binding to the same basic groove path on the spike, preventing binding to HS while exerting mechanistic and allosteric effects that block the interaction with ACE2 and the subsequent entry ^18^. However, the use of heparin as a SARS-CoV-2 antiviral has largely been restricted because its anticoagulant properties can cause adverse side effects in COVID-19 patients. (see Tab.S1). Therefore, chemically modified heparin analogs with reduced anticoagulant effects offer a promising strategy for improving COVID-19 outcomes.

K5 polysaccharides produced by *E. coli* share the same structure as the biosynthetic precursor of heparin/HS and can be chemically or enzymatically modified to resemble heparin or HS structures ^24^. Sulfated K5 derivatives have anti-inflammatory and anti-adhesive effects, no cell toxicity, reduced anti-prothrombin time (APTT) and, a negligible anti-coagulant effect compared to heparin as assessed by an anti-factor Xa assay ^25, 26^. Due to their better pharmacological properties, K5 polysaccharides present an attractive option for antiviral agents. Chemical modifications aim at tailoring their sulfation degree and pattern, including N-deacetylation/N-sulfation and/or O-sulfation, while enzymatic modifications are required for the conversion of GlcA into IdoA (C5 epimerization) ^24, 27, 28^. Non-epimerized K5 derivatives resemble HS structures, while K5 epimerized compounds have a heparin-like structure ^24^. Previous studies showed that sulfated K5s inhibit the infection of a variety of viruses that utilize HSPGs as receptors for cell entry by mimicking the binding interactions of HS, thereby competitively blocking viral access to these host-cell co-receptors ^24^. The antiviral effect of sulfated K5 derivatives on SARS-CoV-2 remains to be investigated.

In this work, we leveraged atomic-detail knowledge of the mechanistic effects of heparin and HS on spike to guide the screening of non-epimerized, variably sulfated K5 derivatives against the SARS-CoV-2 Wuhan-Hu-1 and Omicron BA.1 variants by applying a multidisciplinary approach combining computational molecular modeling and simulation with biochemical and cell-based assays (Fig.S1).

## Results and Discussion

The atomic-level insights obtained from our simulations of spike-heparin complexes^18^ guided the selection of variably sulfated K5 derivatives as potential antiviral candidates targeting the SARS-CoV-2 spike. Specifically, non-epimerized variably sulfated K5 derivatives lacking the active pentasaccharide required for ATIII-dependent anti-coagulant effect were engineered to compete with HS for binding along the spike basic groove while preserving a sulfation pattern capable of inducing structural rearrangements that hinder ACE2 binding. An unsulfated K5 derivative was used as a negative control based on atomic-level insights that indicated that it would not bind to the spike due to the lack of sulfation groups. This has also been used as a control for other heparin-binding proteins^25^.

### K5OSH and K5NOSH bind to spike and exhibit higher activity than heparin

Preliminary screening of a library of K5 derivatives was conducted using biochemical assays to accelerate their evaluation and selection. K5 derivatives (Fig.1A, Tab.S2) were assessed for their ability to bind the Wuhan-Hu-1 spike using a microscale thermophoresis (MST) assay ^29^. All the sulfated K5 derivatives tested and heparin, but not the parental unsulfated K5 (K5), bind to spike in the preliminary binding check procedure (Fig.S2). Unsulfated K5 was excluded from subsequent biochemical assays as it did not bind to spike (Fig.S2). Dose-dependent assays performed on spike-binding GAGs showed that K5 with a high degree of sulfation in O positions (K5OSH) or both N and O positions (K5NOSH) have the highest binding affinity for spike, with dissociation constants (K_d_) of 53.1±7.4 and 36.0±9.8 μM, respectively. K5 with a low degree of sulfation at O positions (K5OSL) or both N and O positions (K5NOSL), N-sulfated K5 (K5NS), and heparin have lower binding affinities, with K_d_ values ranging from 0.2 to 1.5 mM (Fig.1B, Tab.1).

**Fig. 1.**
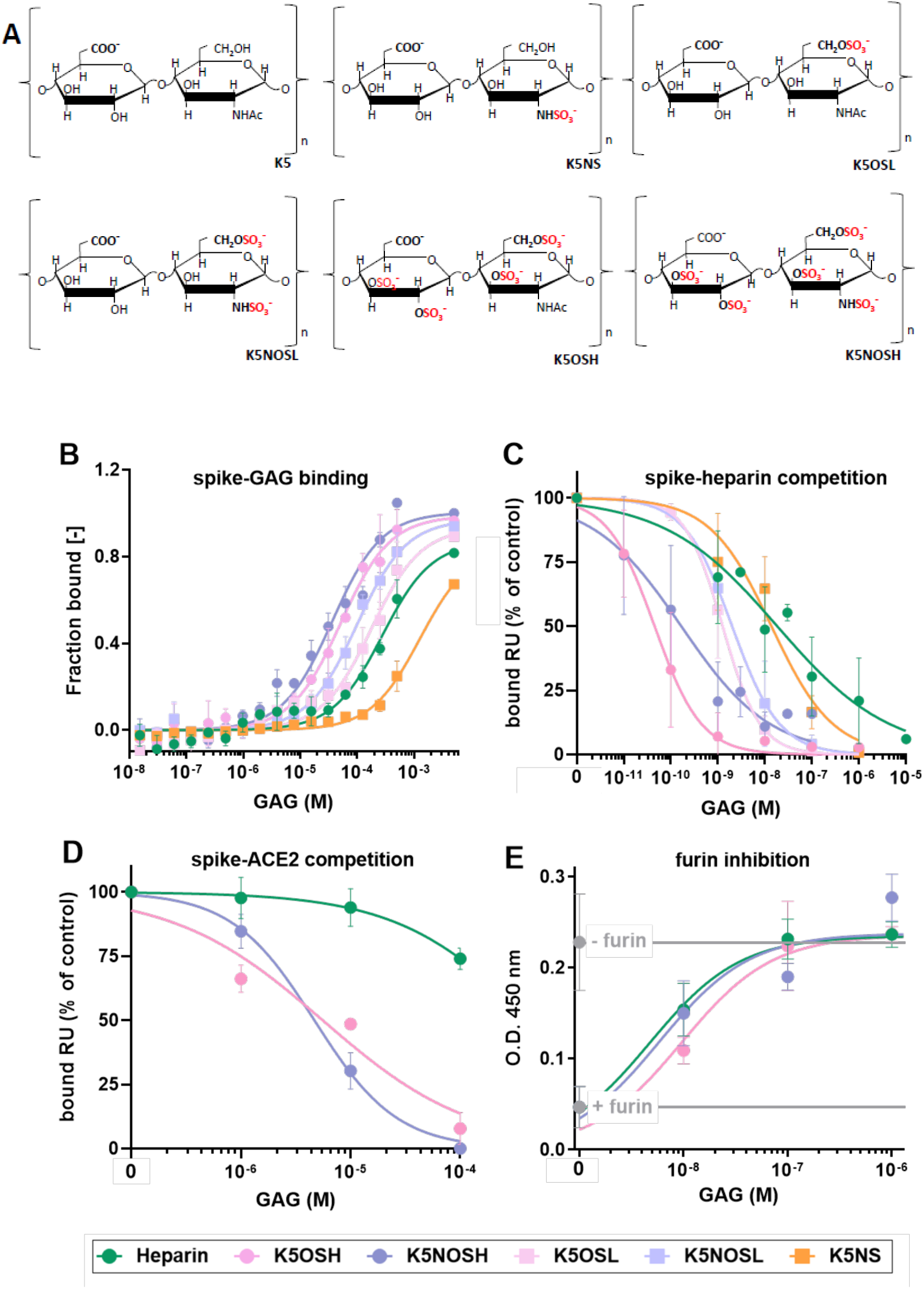
Measurements of effects of sulfated K5 derivatives on spike show K5OSH and K5NOSH outperform heparin in binding and inhibition. **(A)** Predominant composition of the disaccharide unit present in each K5 derivative (ranging from 70% to 100% of the total sequence) is shown (see Tab.S2 for further details). Disaccharides are represented in their ionic form (sodium salt is the common counterion). **(B)** Fraction of spike bound to GAG at increasing concentrations of the indicated GAGs as measured by MST. **(C)** Inhibition of the interaction of immobilized heparin with spike by increasing concentrations of the indicated GAGs as measured by SPR. The responses are plotted as a percentage of the binding of spike measured in the absence of free GAG. **(D)** Inhibition of the binding of spike RBD to immobilized-ACE2 at increasing concentrations of the indicated GAGs as measured by SPR. The responses are plotted as a percentage of the binding of spike measured in the absence of free GAG. **(E)** Inhibition of cleavage of a peptide fragment containing the S_1_/S_2_ basic domain of spike by increasing concentrations of the indicated GAGs. The peptide was left untreated (−furin) or exposed to furin (25 ng/well) (+ furin) (gray points and lanes) and after incubation in the absence and the presence of the GAG, spike cleavage was evaluated by optical density (O.D.) measurements as described in Experimental Procedures. –furin *vs* + furin: P value < 0.00001. Each point is the mean ± sd of three to ten separate determinations (see Tab.1 for more details).

**Tab. 1.** Comparison of measured binding affinity and IC50 values of heparin and the K5 derivatives. The number of replica experiments are reported in brackets. Student’s *t*-test, K5 derivatives *vs* heparin: * P < 0.05; ** P< 0.005. *** P < 0.0005. nd stands for not determined and it means that it was measured but a value was not computed.

| GAG | K<sub>d</sub> ± sd (μM) | IC<sub>50</sub> ± s.d., (nM) |  |  |
| --- | --- | --- | --- | --- |
|  | MST assay: spike affinity binding | SPR assay: spike-heparin interaction inhibition | SPR assay: spike-ACE2 interaction inhibition | Colorimetric assay: spike cleavage by furin |
| Heparin | 355.1±187 (4) | 24.5±18,7 (4) | >100,000 (3) | 8.5±2.1 (4) |
| K5NS | 1,569±800 (3) * | 14.4±4.3 (3) | nd | nd |
| K5OSL | 204.7±75.8 (3) * | 3.5±1.6 (3) | nd | nd |
| K5NOSL | 1,059±10.3 (5) ** | 2.3±1.2 (3) | nd | 11.8±2.4 (3) |
| K5OSH | 53.1±7.4 (3) * | 0.09±0.05 (4) * | 6,300±2,200 (3) *** | 9.9±6.62 (4) |
| K5NOSH | 36.0±9.8 (3) * | 1.1±0.7 (7) ** | 4,700±970 (3) *** | 8.5±2.1 (4) |

A surface plasmon resonance (SPR) competitive binding assay was performed to test the ability of sulfated K5 to prevent spike/HS binding. The HS-analogue heparin was immobilized on an SPR biosensor. All sulfated K5 derivatives tested inhibit in a dose-dependent manner the binding of spike to surface-immobilized heparin (Fig.1C). Heparin and K5NS exhibit modest inhibitory potency (IC_50_ = 29.3±15.5 and 22.3±11.4 nM, respectively), while K5OSL and K5NOSL have significantly higher inhibitory potency (IC_50_ = 1.40±1.0 and 2.65±1.60 nM, respectively). K5OSH and K5NOSH are the strongest heparin antagonists (IC_50_ = 0.2-0.3 nM range) (Tab.1). Next, the ability of K5OSH and K5NOSH to inhibit the binding of the spike RBD to ACE2 immobilized onto a SPR biosensor was examined. In these experimental conditions, heparin poorly inhibits spike RBD-ACE2 binding (IC_50_ > 100,000 nM), whereas K5NOSH and K5OSH exhibit a stronger dose-dependent inhibition with IC_50_ values equal to 4,700±970 and 6,300±2,200 nM, respectively (Fig.1D and Tab.1). Despite previous studies suggesting a direct binding of heparin to ACE2 ^15^, neither heparin nor the sulfated K5 derivatives bound to ACE2 when injected onto the biosensor (Fig.S3).

Spike cleavage at the S1/S2 site mainly occurs in the Golgi apparatus. Nevertheless, this intracellular cleavage can be partial, requiring extracellular furin to cleave the remaining virions ^3, 8, 11, 12, 18, 30^. Heparin inhibits furin cleavage of the spike S1/S2 site *in vitro*^18^. In the same experimental conditions, K5NOSH and K5OSH exhibit similar potency to heparin (Fig.1E and Tab.1).

Collectively, these data indicate that K5OSH and K5NOSH could act as HS antagonists with higher activity than heparin. Consequently, we focused on investigating the antiviral effect of these two promising K5 derivatives.

### K5OSH and K5NOSH act on the spike through direct and allosteric effects

We carried out a total of 18µs of molecular dynamics (MD) simulations of four systems consisting of models of the Wuhan-Hu-1 spike fully glycosylated in either the closed-inactive or the open-active conformation in complex with three K5OSH or K5NOSH chains (Fig.S4). Simulations of the spike in both open and closed conformations were performed to reflect the mixture of these states present in the batches used for the biochemical assays as well as on the SARS-CoV-2 virion. The results were compared with those of our previous simulations of spike-heparin complexes ^18^.

All simulated systems converged to stable structural ensembles within ∼400ns, as shown by the root mean square deviation (RMSD) relative to each starting structure (Fig.S5-S8). Compared to the complex with heparin, the subunits of the closed spike with K5OSH (Fig.S5) and of the open spike with K5NOSH (Fig.S7) have approximately 1 and 2 Å higher RMSD, respectively. In agreement, the root mean square fluctuations (RMSF) show increased values for the NTD and RBD of the closed-spike subunits (Fig.S9) and for the up-RBD of the open-spike complexed upon binding of both K5 derivatives (Fig.S10). The K5OSH and K5NOSH chains maintain H-bonding interactions primarily with the cationic residues of the NTD and S1/S2 of one spike subunit (subunit *n*) and with cationic residues in the RBD of the adjacent spike subunit (subunit *n+1*) with either inactive or active spike bound (Fig.2A-B and 3A-B and Tab.S3-S5). In the closed spike models, the residues involved in the binding of the two K5 derivatives are conserved and similar to those involved in binding heparin (Tab.S3-4). In the open spike models, K5OSH and K5NOSH lie along the basic groove path of the active spike, although interactions with residues in the NTD and RBD significantly differ compared to heparin (Tab.S3, S5).

**Figure 2.**
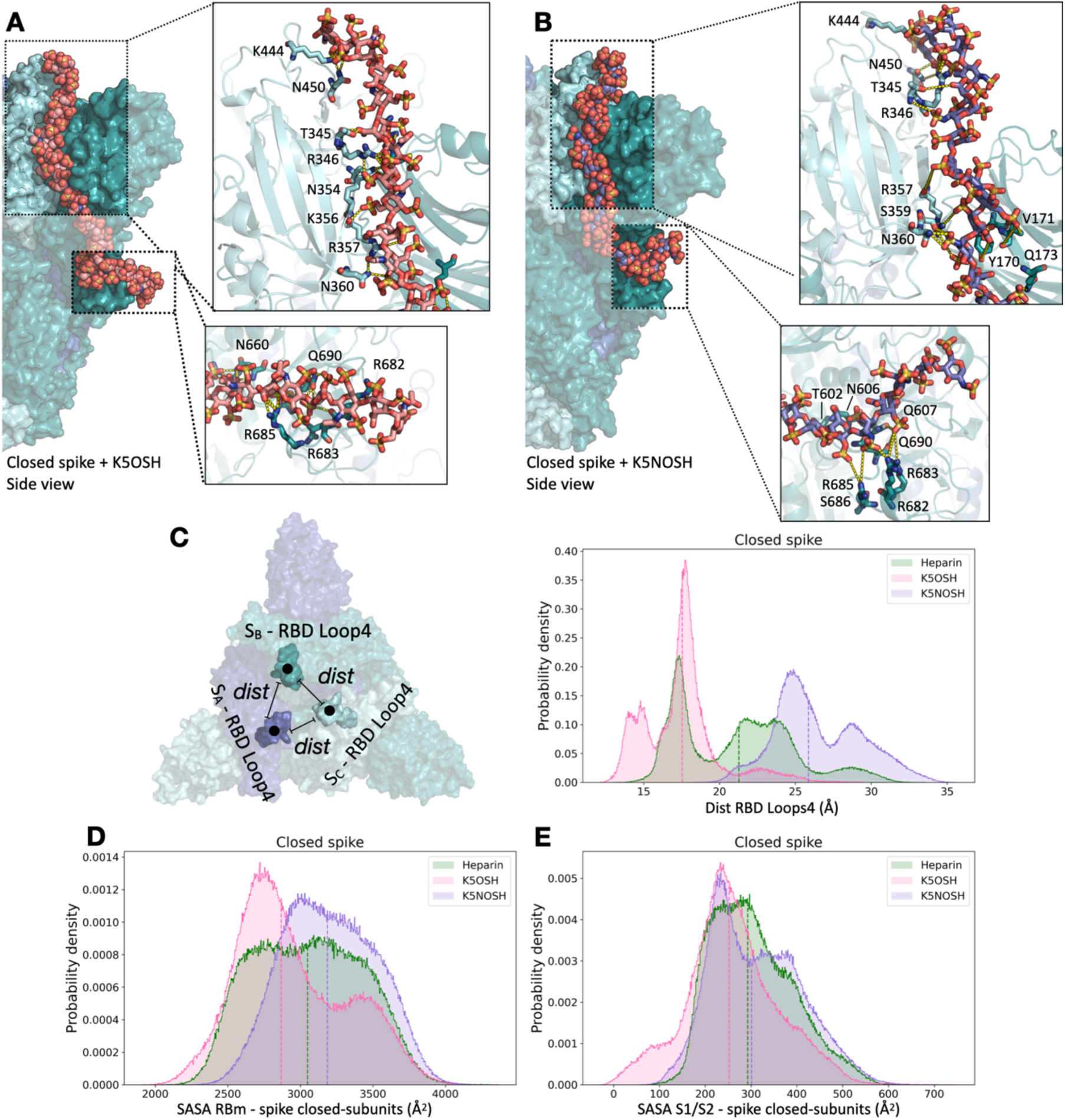
Molecular dynamics simulations of K5OSH and K5NOSH polysaccharides on the closed spike conformation reveal tighter packing of the spike subunits when K5OSH rather than heparin is bound. **(A, B)** Representative structures obtained after MD simulation of the spike bound to three K5OSH **(A)** or K5NOSH **(B)** chains. The S_A_, S_B,_ and S_C_ spike subunits are shown as molecular surfaces in cyan, teal, and blue, respectively. The 31mer K5 chains that span from the RBD of one spike to the S1/S2 site of the adjacent subunit are shown in van der Waals sphere representation colored by element with carbons in pink and purple for K5OSH and K5NOSH, respectively. Insets show the interactions of the K5 chains with residues in the down-RBD (top) and S1/S2 sites (bottom). The spike and its interacting residues are depicted in cartoon and stick representations, respectively, and colored according to the respective subunit. H-bonds are shown as yellow dashed lines. N-glycans covalently attached to the spike are not shown for clarity. **(C)** To define the variation in packing of the closed spike, the distances (dist) between RBD Loop4 (centroids of residues 495-516 represented as spheres) of adjacent spike subunits were calculated. The probability density of the distances between RBD Loop4s was calculated from the converged MD trajectories for all the replicas and colored according to systems: heparin-bound (heparin in green, K5OSH in pink, K5NOSH in purple). K5OSH binding promotes a tighter packing of the closed spike. The probability distribution of the SASA for the down-RBm **(D)** and S1/S2 multifunctional domain **(E)** were calculated from the converged MD trajectories for all the replicas and colored according to systems: heparin-bound (green), K5OSH-bound (pink) and K5NOSH-bound (purple). K5OSH binding reduces the exposure of the RBm and S1/S2 with respect to the reference systems with heparin. Corresponding plots, shown as a function of time and as probability density for each replica trajectory and reference system for **(C-E)** are shown in Fig.S11-13.

In closed spike simulations, the distances between the RBD-Loop4 loops (centroids of residues 495-516) of two adjacent spike subunits were monitored along the trajectories. The binding of K5OSH significantly reduces while K5NOSH increases the average distance between the centroids compared to heparin (Fig.2C and Fig.S11). The absence of direct contact between the RBD-Loop4 and the K5 polysaccharides indicates that they allosterically affect the motion of this highly flexible region of spike RBD (Tab. S3-4). The solvent-accessible surface area (SASA) of spike down-RBDs is significantly reduced only upon K5OSH binding (Fig.2D and Fig.S12). Similarly, SASA of S1/S2 is significantly reduced upon K5OSH binding while K5NOSH binding has a comparable effect to heparin (Fig.2E and Fig.S13).

In open spike simulations, the K5 derivatives affect the movement of the spike up-RBD (Fig.3A-B). The RMSD of residues 527-PKK-529, the hinge-residues responsible for the RBD movement ^18^, reveals stabilization of the residues of the up-RBD with K5OSH bound while K5NOSH derivatives have a comparable effect to heparin (Fig.3C and Fig.S14). Dihedral Principal Component Analysis (dPCA) highlights an altered sampling of the conformational space of the up-RBD hinge residues upon binding of K5 derivatives (Fig.S15). In the presence of K5OSH, the first eigenvector describing the up-RBD motion showed overall stabilization of the domain, pointing to a potential inhibitory gating effect on the host cell-receptor binding (Fig.3D and Fig.S16). With K5NOSH, the first eigenvector reveals that the polyanionic compounds promote the transition of the RBD from the starting “up” state to a semi-closed/closed state (Fig.3D and Fig.S16) leading to greater shielding of the up-RBm than heparin or K5OSH (Fig.3E and Fig.S17). K5OSH derivatives have an effect comparable to heparin in shielding the S1/S2 functional site of the spike up-subunit while K5NOSH derivatives have a significantly lower ability to mask this functional site (Fig.3F, Fig.S18 and Tab.S3). Notably, both K5s outperform heparin in shielding the S1/S2 site of the closed subunits (S_A_ and S_B_) on the open spike (Fig.S18).

**Figure 3.**
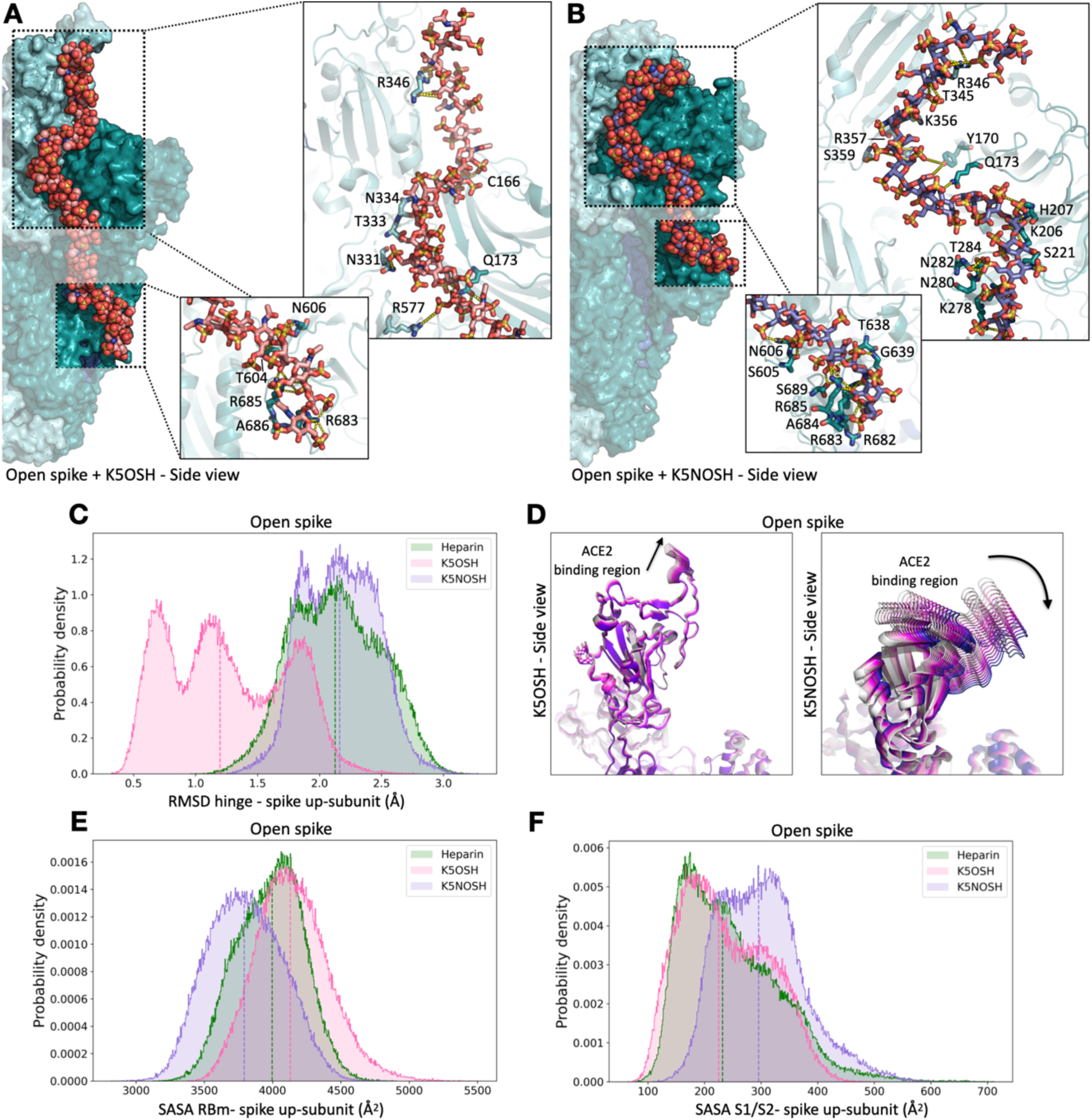
Molecular dynamics simulations of K5OSH and K5NOSH polysaccharides on the open spike conformation show that, compared to heparin, K5OSH affects the RBD-orientation while K5NOSH alters exposure of spike binding sites. **(A,B)** Representative structures obtained after MD simulation of spike bound to three **(A)** K5OSH and **(B)** K5NOSH chains. The S_A_, S_B,_ and S_C_ spike subunits are shown as surface in cyan, teal, and blue, respectively. The 31mer K5 chains that span from the RBD of one spike to the S1/S2 site of the adjacent subunit are shown in sphere representation colored by element with pink and purple representation for K5OSH and K5NOSH, respectively. Insets show the interactions between the K5 polysaccharides and residues in the up-RBD (top) or S1/S2 sites (bottom). The spike and its interacting residues are depicted in cartoon and stick representations, respectively, and colored according to the subunit to which they are attached. H-bonds are shown by yellow dashed lines. N-glycans covalently attached to the spike are not shown for clarity. **(C)** The probability density of the RMSD for the hinge region (residues 527-529) of the spike up-subunit calculated from the converged MD trajectories for all the replicas and colored according to systems (heparin in green, K5OSH in pink, K5NOSH in purple) showing that K5OSH-binding results in stabilization of the hinge region in the spike up-RBD. **(D)** The change in motion of the up-RBD induced by the presence of K5 derivatives is shown by the superimposition of conformations extracted at equal time intervals along the trajectories (from magenta to blue) and projected onto the first essential dynamics eigenvector with K5OSH (left) and K5NOSH (right). The RBD is shown in cartoon representation and the K5 derivatives are omitted for ease of visualization. The probability distribution of the SASA for the up-RBD RBm (**E**) and S1/S2 multifunctional domain (**F**) calculated from the converged MD trajectories for all the replicas and colored according to systems (heparin in green, K5OSH in pink, K5NOSH in purple) showing that only K5NOSH decreases the exposure of the RBD RBm whereas K5OSH has a comparable effect to heparin in shielding the exposure of the S1/S2 basic site of the up-spike subunit. Corresponding plots, shown as a function of time and as probability density for each replica trajectory and reference system for **(C-E)** are shown in Fig.S14-18.

In summary, the K5OSH derivative outperforms heparin and K5NOSH as regards spike inhibition by promoting a more compact packing and shielding of closed spike while promoting rigidification of the up-RBD in the open spike thereby hampering the binding to ACE2 and enhancing the shielding of S1/S2. K5NOSH shields the RBm of the spike up-subunit better than heparin and K5OSH, and thus hampers ACE2 binding. Computational studies provide a mechanistic explanation of the stronger activity of the K5 derivatives than heparin in biochemical assays while pointing to K5OSH as the most promising among the compounds tested.

### K5OSH and K5NOSH inhibit SARS-CoV-2 infection by blocking S-mediated membrane fusion

Biochemical and computational assays point to the ability of K5OSH and K5NOSH to exert an antiviral effect by multiple mechanisms including competing with the host-cell HSPGs for the binding of spike, preventing the spike-proteolytic priming at the S1/S2 site, by directly shielding it, and at the S2’ site by hampering the binding of spike to ACE2 and the subsequent exposure of residue R815 ^13, 17^.

To gain further insight into the mechanism of the inhibition, the effect of the K5 compounds was first tested directly on spike-mediated membrane fusion using a syncitia assay, which depends on the S2’ cleavage carried out by TMPRSS2 at the plasma membrane ^31^. The assay was performed on cells expressing either Wuhan-Hu-1 spike, corresponding to the biochemical and computational assays, or Omicron BA.1 spike, to evaluate the therapeutic potential of the K5 derivatives on a more recent virus variant. Using automated microscopy and machine learning-based segmentation of nuclei in spike-positive cells and spike-positive syncitia, we determined the fraction of nuclei involved in syncytia formation. Our data show that the number of Wuhan-Hu-1 spike-positive cells involved in syncytia formation using VeroE6 cells is higher than the number of Omicron BA.1 spike-positive cells (Fig.4A-B). Syncytia contained fewer nuclei upon treatment with K5 derivatives (Fig. 4C-D). K5OSH had a statistically significant inhibitory effect on Wuhan-Hu-1 induced fusion while both K5OSH and K5NOSH significantly reduced Omicron BA.1 membrane fusion.

**Figure 4.**
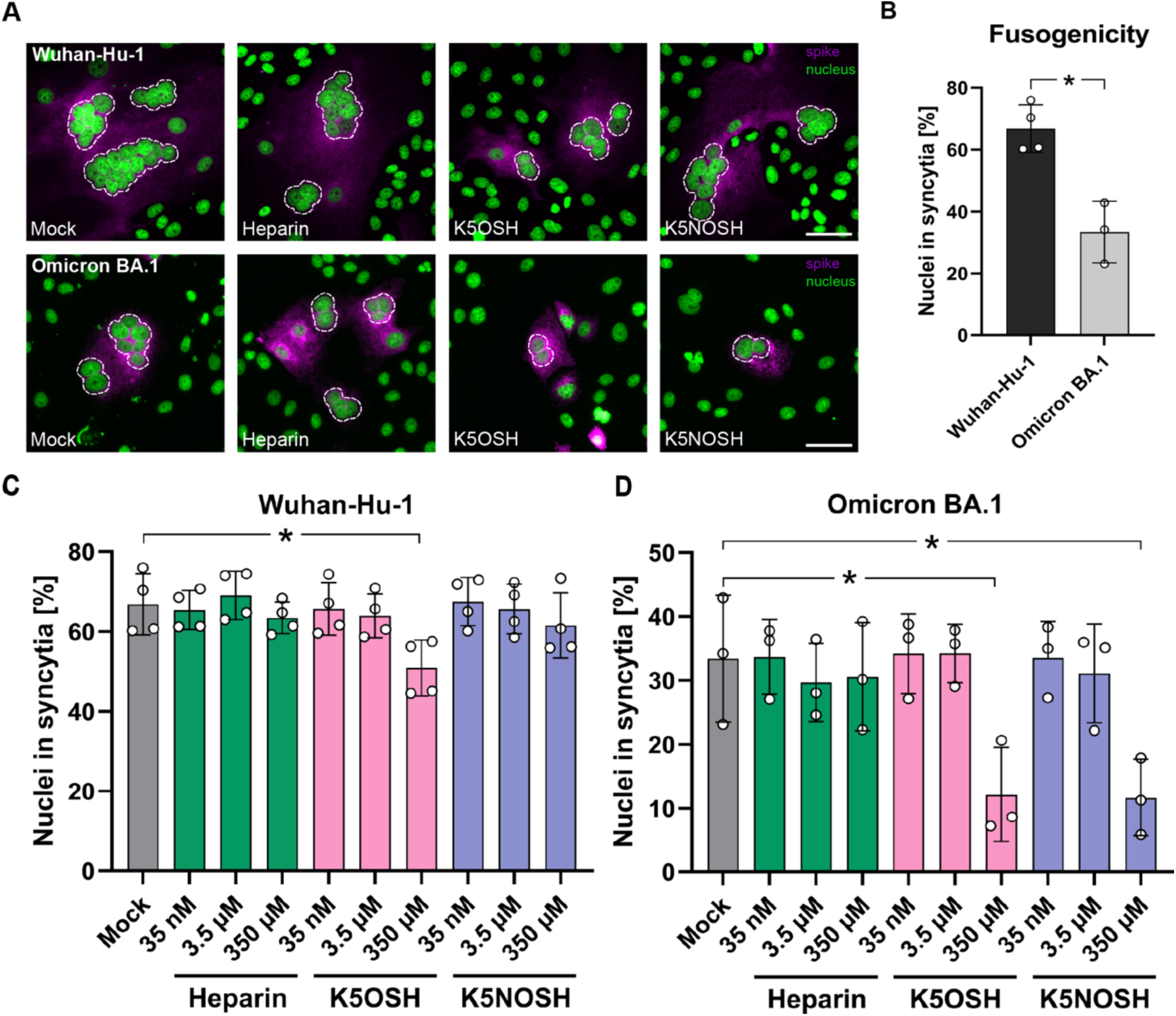
Measurements of syncytia formation show greater dose-dependent effects of K5 compounds than heparin. (A) Representative images of syncytia formation assay in VeroE6 cells upon treatment (350 µM) with K5 compounds. Scale bar: 50 µm. (B) Number of nuclei involved in syncytia formation is higher in Wuhan-Hu-1 spike-positive cells than in Omicron BA.1 spike-positive cells. (C) Effect of K5 compounds on syncytia formation induced by Wuhan-Hu-1 spike. (D) Effect of K5 compounds on syncytia formation induced by Omicron BA.1 spike. Only spike-positive cells were quantified. Data of four (Wuhan-Hu-1) and three (Omicron BA.1) independent experiments. Values are mean with SD. P values determined by Welch’s t-test: * P < 0.05; ** P< 0.005.

Finally, K5OSH and K5NOSH were examined for their antiviral activity in an infection assay using authentic SARS-CoV-2 B.1 (D614G) and Omicron BA.1 variants on VeroE6 and on A549 ACE2+ cells. Unsulfated K5 was used as negative control (Fig.S2). Cell viability assays were conducted in the absence of the virus to calculate the half-cytotoxic concentration (CC_50_) for the compounds. At 48 h post-treatment, all compounds exhibited no significant toxicity at concentrations up to 100 μM on VeroE6 cells (Fig.5A). Heparin and K5OSH showed a slight cytotoxic effect at 100 μM on A549 ACE+ cells, and therefore this dose concentration was excluded from the subsequent assays (Fig.5B). Unsulfated K5 was ineffective against the B.1 and BA.1 variants on both VeroE6 and A549 ACE+ cells (Fig.5C-F). When tested against the B.1 variant, K5OSH and K5NOSH had a weak inhibitory effect in VeroE6 cells (IC_50_ = 17.99 and >10 μM, respectively) (Fig.5C and Tab.2). In A549 ACE+ cells, K5OSH exhibited stronger antiviral activity (IC_50_ = 0.2 µM) than K5NOSH and heparin (IC_50_ = 2.9 and 2.55μM, respectively) (Fig. 5D and Tab.2). When tested against the Omicron BA.1 variant, K5OSH and K5NOSH inhibited infection in both VeroE6 cells (IC_50_ = 0.4 and 8.0 μM, respectively) and A549 ACE+ cells (IC_50_ = 1.9 and 3.8 μM, respectively) while heparin was ineffective (Fig. 5E-F and Tab.2).

**Tab. 2.** Inhibition of SARS-CoV-2 infection in cells by heparin, K5, K5OSH, and K5NOSH. In the absence of a saturation effect, the IC_50_ values were calculated on the trend of the curve (Fig.5) using GraphPrism 8.4. The number of repeated experiments is reported in brackets. Data of two independent experiments.

| Inhibitory potency (IC<sub>50</sub> ± sd, μM) |  |  |  |  |
| --- | --- | --- | --- | --- |
|  | VeroE6 cells |  | A549 ACE2+ cells |  |
| GAG | B.1 | BA.1 | B.1 | BA.1 |
| Heparin | 537.4 | 160.1 | 2.55 | 25.16 |
| K5 | > 10 | > 10 | > 10 | > 10 |
| K5OSH | 17.99 | 0.4 | 0.2 | 1.9 |
| K5NOSH | > 10 | 8.0 | 2.9 | 3.8 |

**Figure 5.**
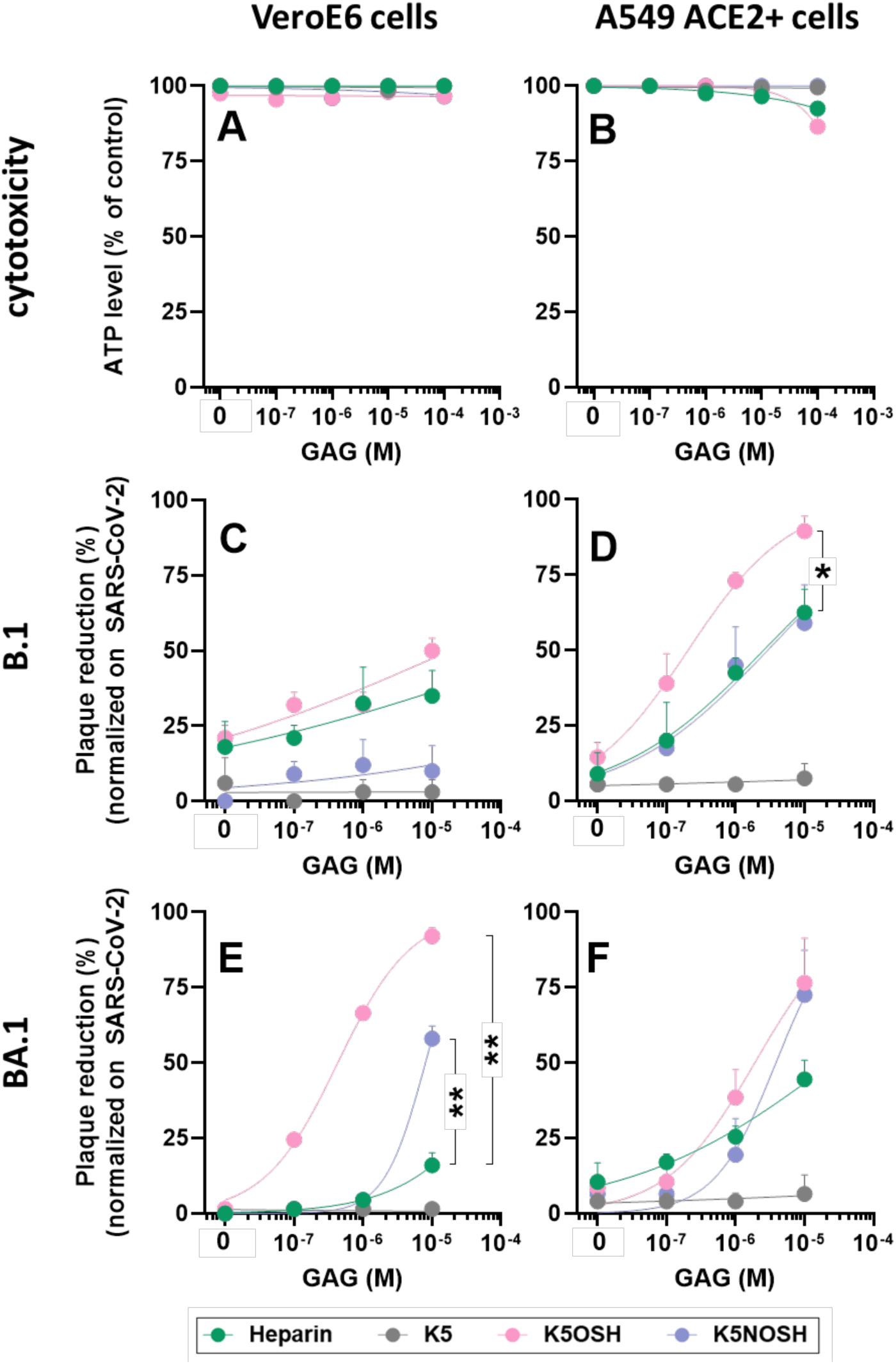
Measurements of cytotoxicity and antiviral activity against SARS-CoV-2 B.1 and Omicron BA.1 show inhibition by the K5 compounds with K5OSH showing the greatest inhibitory effect. VeroE6 cells or A549 ACE2+ cells were treated with increasing concentrations of heparin, K5, K5OSH, and K5NOSH. **(A, B)** The cells remained viable in the presence of heparin and the K5 compounds as evaluated by measuring the ATP levels. VeroE6 or A549 ACE2+ cells were infected with the B.1 **(C, D)** or Omicron BA.1 **(E, F)** isolates in the presence or the absence of increasing concentrations of heparin, K5, K5OSH, and K5NOSH. Infection was reduced in a concentration-dependent manner as shown by the percentage of plaque reduction compared to SARS-CoV-2 alone. Data are presented as the mean value ± standard error of two independent replicates. * P < 0.05; ** P< 0.005.

Collectively, our results highlight the higher antiviral activity of K5OSH and K5NOSH compared to heparin against SARS-CoV-2, and particularly against the Omicron variant.

## Discussion

Heparin has been investigated for its antiviral effect against SARS-CoV-2 ^21–23^. It competes with HS for binding to the basic groove path along the spike and thereby directly and allosterically hinders the binding to the host-cell receptor ACE2, spike proteolytic cleavage, and its subsequent activation ^18^. However, the anticoagulant activity of heparin has hampered its application as an antiviral agent. K5 polysaccharides derived from *E. Coli* share the same structure as the HS/heparin precursors while exerting anti-inflammatory and anti-adhesive effects, and having no cell toxicity, reduced anti-prothrombin time (APTT), and negligible anti-Xa effects ^25, 26^.

Here, by leveraging the atomic-detail knowledge obtained by simulating spike-heparin and spike-HS-ACE2 complexes ^17, 18^, we screened a library of non-epimerized variably sulfated K5 derivatives, including an unsulfated K5 derivative used as a negative control, aimed at targeting the spike basic groove path that accommodates heparin/HS and competitively preventing their binding while simultaneously hindering ACE2 binding and spike proteolytic cleavage.

First, a screening based on biochemical assays revealed that increasing the sulfation degree of the K5 derivatives significantly improved their affinity for the Wuhan-Hu-1 spike. K5OSH and K5NOSH are the strongest binders with K_d_ values 6-10 times lower than heparin. Hence, we focused on these compounds. K5OSH and K5NOSH inhibit the cleavage of the spike S1/S2 site by furin and the binding of Wuhan-Hu-1 spike-RBD to HS-analogue heparin and ACE2.

Next, computational studies demonstrated that K5OSH and K5NOSH bind to the Wuhan-Hu-1 spike following the basic groove previously identified, and competing with host-cell HSPGs for binding while shielding the S1/S2 functional site, therefore preventing furin cleavage ^17, 18^. Strong electrostatic repulsion between K5OSH or K5NOSH (both with a net charge of −75e for a 31mer) and the ACE2 receptor binding domain (−27e), coupled with shielding of the spike RBm, inhibit the binding of the host cell receptor ACE2, thereby hampering the subsequent cleavage of S2’ by TMPRSS2 ^13, 17^. Mechanistically, K5OSH derivatives outperform heparin and K5NOSH, promoting a better packing of the closed spike and rigidification of the up-RBD of the open spike, pointing to a potential inhibitory gating effect on host cell-receptor binding.

Subsequently, an assay of syncytia, which depend on the S2’ cleavage carried out by TMPRSS2 at the plasma membrane, was performed. It has been shown that syncytia formation is enhanced by TMPRSS2 and inhibited by interferon-induced transmembrane protein 1 ^31^. SARS-CoV-2 variants show differences in the preferred route of entry and the site of membrane fusion. The original Wuhan-Hu-1, B.1 and Delta variants are highly sensitive to Cemostat (an inhibitor of TMRSS2) and prefer to enter the cell by direct fusion with the plasma membrane ^32^. In contrast, the Omicron variants are less dependent on TMPRSS2 cleavage and utilize an endocytic pathway ^33, 34^ where both S1/S2 and S2’ cleavage can be carried out by endosome-associated proteases ^35, 36^. Hence, the effect of K5 compounds on spikes of the Wuhan-Hu-1 and Omicron BA.1 variants was evaluated using syncytia assays in VeroE6 cells, which contain low amounts of TMPRSS2 ^36^. In agreement with previous reports, the expression of the Wuhan-Hu-1 spike induces more cells to form syncytia than the expression of the Omicron BA.1 variant spike protein ^33^. Notably, K5OSH significantly inhibits syncytia formation induced by the Wuhan-Hu-1 spike protein. K5OSH and K5NOSH both significantly inhibit Omicron BA.1 spike-mediated VeroE6 syncytia formation at the highest used concentration. These results indicate that the K5 inhibitory mechanism is linked to the inhibition of proteolytic cleavage at the S1/S2 and S2’ cleavage sites, as indicated in the computational studies.

Finally, K5OSH and K5NOSH were evaluated for their capacity to inhibit infection by SARS-CoV-2 B.1 and Omicron BA.1 variants in VeroE6 and A549 ACE2+ cells (both expressing low levels of TMPRSS2). The K5 derivatives tested exerted a stronger inhibitory activity on infection by Omicron BA.1 and K5OSH was more potent than K5NOSH at inhibiting virus infection, in line with syncytia formation assays. One explanation of the different activity of the K5 derivatives in infection assays using alternative cell types can be attributed to variations in the ratio of closed to open spikes on SARS-CoV-2 virions. In cells with low ACE2 expression, such as Vero E6 cells, spikes predominantly adopt the prefusion closed conformation ^2^. Computational studies indicate that in closed spike conformation, K5OSH exhibits a higher antiviral effect compared to the other polysaccharides investigated. In contrast, the ratio between open and closed spikes is balanced in cells with higher ACE2 expression as A549 ACE2+ cells ^2^. This balance explains the increased efficacy of heparin and K5NOSH, as computational studies attribute their enhanced activity to a greater effect on the open spike, although K5OSH derivatives remain the most active overall. Furthermore, since the interaction between K5 and the spike is driven by long-range electrostatic interactions, the enhanced activity of K5 derivatives against the Omicron BA.1 variant is likely due to its increased net charge compared to the Wuhan-Hu-1 spike,^17, 20^, resulting in greater affinity for the selected K5 derivatives.

Overall, the multidisciplinary approach applied here demonstrates that K5OSH is the most potent anti-SARS-CoV-2 agent out of the tested compounds. It competes with HS for spike binding, inhibiting its proteolytic priming at S1/S2 by direct shielding, and at S2’ by blocking ACE2 binding, the subsequent exposure of R815, and TMPRSS2 cleavage ^13, 17^. Of note, K5OSH derivatives are not susceptible to heparinase I and II degradation due to their specific sulfation pattern ^37^. Nevertheless, given the increased levels of cellular lyases in the plasma of COVID-19 patients with severe clinical outcomes, further experiments aimed at excluding the susceptibility of K5OSH to cell lyases would be crucial. Furthermore, since the effects of the selected K5 derivative have been tested on only two spike variants, further experiments are needed to elucidate antiviral potential and impact on infectivity and inhibitor binding on circulating SARS-CoV-2 variants and cells overexpressing TMPRSS2. Finally, the biosafety and effectiveness of these compounds need a more detailed investigation and evaluation using animal models before pharmaceutical applications.

SARS-CoV-2 has evolved to target the upper respiratory tract. Inhaled nebulized heparin was tested to treat COVID-19 patients but its bioavailability in the blood of patients has hampered a broader application as an antiviral due to the risk of bleeding or hemorrhagic events ^38–40^. Inhaled nebulized drugs based on sulfated K5 would provide higher drug concentrations topically, resulting in a rapid clinical response, decreased risks of K5 degradation due to cellular lyase, and more importantly, prevention of systemic side effects due to the safer profile of K5 compared to heparin ^25, 26^. Of note, topical administration of sulfated K5 derivatives showed no toxicity and no significant inflammation ^41^. These factors taken together underline the need for further clinical investigations to test the ability of nebulized K5OSH, identified as the most promising candidate in this study, to prevent SARS-CoV-2 infection.

## Materials and Methods

### Reagents

Reagents and materials were used as received, unless otherwise mentioned, and were purchased from the following: Human recombinant SARS-CoV-2 Wuhan-Hu-1 spike His-Tag protein and RBD from Sino Biological (#40592-V08B); ACE2 from Acrobiosystem (#AC2-H52H8); Bovine Serum Albumin (BSA) from Merck (#810037); Human recombinant furin from OriGene Technologies Inc. (#TP304279M); Conventional heparin (13.6 kDa - purity ≥95%) from a commercial batch of unfractionated sodium heparin from Laboratori Derivati Organici S.p.A. (#9041-08-1); Capsular K5 polysaccharides prepared as described in Oreste et. al ^41^.

### Biochemical studies

#### Microscale thermophoresis (MST) assays

Spike His-tag protein (100 nM protein, molar dye:protein ratio ≈ 1:2) was labeled using the His-Tag Protein Labeling Kit RED-tris-NTA (NanoTemper Technologies) in labeling buffer at room temperature for 30 min in the dark and adjusted to 25 nM with phosphate-buffered saline buffer (PBS) containing 0.05 % Tween 20 (PBST buffer). A two-step procedure was followed with an initial binding measurement performed on the Monolith NT.115 (NanoTemper Technologies) using the MO.Control 2 software. Samples containing spike (20 nM in PBS) in the absence or in the presence of 15 μM of each K5 derivative were prepared, incubated for 30 min, centrifuged at 10,000×g for 10 min, and loaded into Monolith NT.115 capillaries (NanoTemper Technologies) using 4 capillaries for each sample. MST was measured using a Monolith NT.115 instrument (NanoTemper Technologies) at 25°C. The instrument parameters were adjusted to 100 % LED power and medium MST power, and the MST signal of spike alone was compared to the MST signal of the complex. The signal-to-noise ratio, defined as the response amplitude (difference of MST signal of spike alone and in complex, in %_o_ F_norm_ units), divided by the noise of the measurement (standard deviation of the 4 replicates) was used to evaluate the quality of the binding data. To confirm the formation of the complex, the signal/noise ratio was required to be higher than 5. Only K5 derivatives that showed binding in this initial step were further analyzed to determine their binding affinity (K_d_) for spike. For this purpose, 16x 1:1 dilutions of each K5 derivative were prepared in PBST buffer. Each dilution was mixed with one volume of labeled spike (final spike concentration equal to 25 nM and final ligand concentrations ranging from 15 nM to 500 μM). After 30 min incubation followed by centrifugation at 10,000×g for 10 min, samples were loaded into capillaries and MST was measured as described. Data from two to three independently pipetted measurements were analyzed using the MO.Affinity Analysis software version 2.3 (NanoTemper Technologies) using the signal from an MST-on time of 5s.

#### Surface plasmon resonance (SPR) assays

A BIAcore X100 instrument (Cytiva) was used. SPR was exploited to measure changes in refractive index caused by the ability of the different sulfated K5 derivatives to prevent the binding of spike to heparin or ACE2 immobilized on a biosensor. Carboxy-methylated dextran CM5 and SA sensorchips, 1-ethyl-3-(3-diaminopropyl)-carbodiimide hydrochloride (EDC) and N-hydroxysuccinimide (NHS) were from Cytiva.

##### Heparin biosensor

The spike-heparin interaction has been already characterized in a SPR setup^16^. The same experimental conditions were adopted here to evaluate the capacity of sulfated K5 derivatives to prevent the binding of spike to surface-immobilized heparin. A research grade sensor chip SA (Cytiva), coated with a carboxy-methylated dextran matrix pre-immobilized with streptavidin, was used. The first flow cell of the sensor chip was conditioned with three consecutive 1-min injections of 1.0 M NaCl in 50 mM NaOH. Then, heparin that had been biotinylated at its reducing end was diluted at 0.015 mg/ml in 10 mM HEPES buffer, pH 7.4, containing 150 mM NaCl, 3 mM EDTA, and 0.005% surfactant P20 (HBS-EP) and injected over the activated surface for 8 min at a flow rate of 5.0 μL/min, allowing the immobilization of 120 resonance units (RU) (equal to 9 fmol/mm^2^) of the GAG. The second flow cell of the sensor chip was coated with biotinylated BSA protein as described above, allowing the immobilization of 569 RU (equal to 9 fmol/mm^2^) of protein, and used to evaluate the nonspecific binding and for blank subtraction. For binding analysis, the different GAGs reported in Tab.S2 were diluted in HBS-EP to concentrations ranging between 0.01 to 10,000 nM with a step size of 10 in the absence or presence of spike protein (45 nM) and injected at a flow rate of 30 μl/min and at a temperature of 20°C. The complex was allowed to associate and dissociate for 120 and 300 s, respectively. At the end of the dissociation phase, the surfaces were regenerated with a 60 s injection of 0.25% SDS.

##### ACE2 biosensor

ACE2 was immobilized on one of the two flow cells of a CM5 sensor chip by standard amine-coupling chemistry as described in Rusnati et al.^42^. BSA was immobilized on the second flow cell which was used for blank subtraction. Different sensor chips were prepared and used for the analyses with the amount of immobilized ACE2 and BSA ranging from 2,400 to 7,300 RU (equal to 19 to 24 femtomoles/mm^2^) and 1,600 to 13,300 RU (equal to 22 to 186 femtomoles/mm^2^) of the two proteins, respectively. For binding analysis, heparin (used as a reference compound) and the K5 derivatives selected from the previous screening performed on the heparin biosensor were diluted in HBS-EP at 1, 10 and 100 µM and injected over the two flow cells in the absence or in the presence of a spike fragment representing its RBD domain (20 nM), as for the heparin biosensor described above. At the end of the dissociation phase, the surfaces were extensively washed until the baseline returned to pre-injection levels.

#### Furin cleavage assay

The ability of the various GAGs to inhibit spike cleavage at the S1/S2 site by furin was evaluated by using the colorimetric assay “CoviDrop^TM^ SARS-CoV-2 targeted proprotein convertase inhibitor screening fast kit” (Epigentek, County Blvd, Farmingdale NY). Briefly, a 2.0 kDa SARS-CoV-2 spike fragment containing the intact S1/S2 _681_RRAR_684_ HBD is tagged with polyhistidine and biotin at its N-terminus and C-terminus, respectively, and immobilized onto microplate wells through histidine-Ni-NTA. The cleavage of the substrate at the S1/S2 site removes the C-terminal S2 portion linked to biotin, causing a decrease of the signal generated by avidin/biotin binding that is detected by colorimetric reaction measured as absorbance at 450 nm in a microplate spectrophotometer). Furin cleavage inhibition blocks the reduction of the signal. Consequently, the extent of spike cleavage is inversely proportional to the signal intensity. The assay was performed according to the manufacturer’s instructions (https://www.epigentek.com/catalog/covidrop-sars-cov-targeted-proprotein-convertase-inhibitor-screening-fast-kit-p-84596.html).

### Computational studies

#### Modeling of the systems

Four different systems were modelled: spike Wuhan-Hu-1 variant fully glycosylated in the closed conformation with (i) three K5OSH or (ii) three K5NOSH chains bound, and in the open conformation with (iii) three K5OSH or (iv) three K5NOSH chains bound, respectively. Atomic coordinates of the complete models of closed and open spike (without the stalk) were retrieved from Paiardi et al.^18^. K5 derivatives of 31 monosaccharides (31mer) were docked onto the spike using the incremental sliding window docking method^43^ and the polysaccharide models reported in Fig.S19. They spanned from the basic domain in the RBD to the S1/S2 functional site. Corresponding models and simulation trajectories of the closed and open conformation of spike with three heparin chains were retrieved from Paiardi et al.^18^ and used for comparison.

#### All-atom molecular dynamics (MD) simulations

Simulations were carried out using Amber20^44^. Spike was parametrized using ff14SB^45^ and GLYCAM-06j^46^ force fields. K5 polysaccharides were parameterized as described previously^43^. All systems were placed in a periodic cubic box solvated using the TIP3P water model^47^ with 10 Å between the solutes and the edges of the box. Na+ and Cl-ions were added to the box to neutralize the systems and immerse them in a solvent with an ionic strength of 150 mM. Four replicas for each system were simulated. Each system was energy minimized in 14 consecutive minimization steps, each of 100 steps of steepest descent followed by 900 steps of conjugate gradient, with decreasing positional restraints from 1000 to 0 kcal/mol A^2^ on all the atoms of the systems, excluding waters, counter-ions, and hydrogens, with a cutoff for non-bonded interactions of 8 Å. The systems were then subjected to two consecutive steps of heating, each of 100,000 steps, from 10 to 100 K, and from 100 to 310 K, in an NVT ensemble with a Langevin thermostat. Bonds involving hydrogen atoms were constrained with the SHAKE algorithm ^48^, and a 2 fs time step was used. The systems were then equilibrated at 310 K in four consecutive steps of 2.5 ns each in the NPT ensemble with a Langevin thermostat with random velocities assigned at the beginning of each step. For each system simulated, four independent replica production runs following the same protocol as for the equilibration were carried out starting from restart files chosen randomly from the last 5 ns of equilibration. During the MD simulations, a cutoff of 8 Å for the evaluation of short-range non-bonded interactions was used, and the Particle Mesh Ewald method was employed for the long-range electrostatic interactions. The temperature was kept constant at 310 K with a Langevin thermostat. Coordinates were written at intervals of 100 ps. Production simulations for replicas of systems (i), (ii) and (iii) were carried out for up to 1 µs, while for system (iv), simulations were extended up to 1.5 µs.

#### Analysis of MD simulations

MD trajectories were analyzed using CPPTRAJ^49^ in AmberTools20^49^, and molecular graphics analysis was performed using Visual Molecular Dynamics (VMD)^50^. Data obtained were then compared with previous analyses performed for simulations of inactive and active spike in the presence of three heparin chains^18^. For consistency, the analysis was performed using the same scripts and parameters as used by Paiardi et al.^18^.

– *Hydrogen bond (H-bond) analysis* was performed using CPPTRAJ^49^ along all the trajectories for frames at intervals of 10 ns (coordinates written every 100 ps collected with a stride of 100 frames) and setting 3.5 Å as the upper distance for defining a H-bond between heavy atoms. All the atoms, including the hydrogens, of the systems were considered (Fig.2A-B, Fig.3A-B, Tab.S3-S5).
– *Distances between RBD-Loop4s* were computed using CPPTRAJ^49^ along all the trajectories. Centroids were calculated for the RBD-Loop4 (residues 495-516) of each spike subunit and the distance between two centroids was computed along the trajectory (Fig.2C, Fig.S11).
– *Solvent-accessible surface area (SASA)* was computed using CPPTRAJ^49^ along the trajectory with a van der Waals radius of the solvent probe of 1.4 Å. For the analysis of the receptor-binding residues, all the residues of the RBD (residues 319–541) were considered along the trajectory (Fig.2D, Fig.3F, Fig.S12, S17) while for the S1/S2-HBD site, residues 682 to 685 were considered (Fig.2E, Fig.3G, Fig.S13, S18).
– *Root mean square deviations (RMSD)* were calculated using CPPTRAJ^49^ for all C-alpha atoms of the individual spike subunits—S_A_, S_B_, S_C_ —and for all the carbon, oxygen, sulfate, and nitrogen atoms of the K5 derivatives (Fig.S5-S8). The RMSDs of the hinge regions were calculated for the C-alpha atoms of residues 527 to 529 for S_A_, S_B_, and S_C_, separately (Fig.3C, Fig.S14).
– *Root mean square fluctuations (RMSF)* were calculated using CPPTRAJ^49^ for all C-alpha atoms of the individual spike subunits—S_A_, S_B_, S_C_—and for all the carbon, oxygen, sulfate, and nitrogen atoms of the K5 derivatives (Fig.S9-S10).
– *Dihedral principal component analysis (dPCA)* was performed using CPPTRAJ^49^. The dihedral covariance matrix and the projection were calculated for the backbone phi/psi angles of residues 527 to 529 of the S_C_ monomer. The first four eigenvectors and eigenvalues were extracted, and the first two principal components were plotted for all of the systems. All the systems were transformed into the same principal component space to evaluate the variance across the replicas (Fig.3D and Fig.S15).
– *Essential dynamics (ED)* analysis was performed using Principal Component Analysis (PCA) of the unbiased MD simulations. PCA was performed along all the trajectories individually with CPPTRAJ^49^. The principal modes of motion were visualized using VMD^50^. The first normalized eigenvectors were plotted along the trajectory, and the direction of motion was defined by visual inspection (Fig.3E and Fig.S16).

### Cells

VeroE6 cells were purchased from the American Type Culture Collection (ATCC; Catalog #CRL-1586) for the syncytia formation assay and from the “Istituto Zooprofilattico Sperimentale della Lombardia e dell’Emilia Romagna” (Brescia, Italy) for infection assays. They were maintained in Dulbecco’s Modified Eagle Medium (DMEM, GlutaMAX supplement, 100 U/ml penicillin, 100 μg/ml streptomycin and 10% fetal bovine serum Gibco, Thermo-Fisher, Waltham, MA, USA) at 37 °C and 5% CO_2_. A549 ACE2-positive (A549 ACE2+) cells, a kind gift from Dr. Stephen J Elledge (Harvard Medical School, Boston, MA, USA), were cultured in RPMI (Gibco, Thermo-Fisher Scientific) supplemented with 10% FBS. Both cell lines were maintained at 37°C in a humidified atmosphere of 5% CO_2_.

### Plasmids and antibodies

Plasmid encoding for SARS-CoV-2 Wuhan-Hu-1 spike (YP_009724390.1) was kindly provided by Prof. Dr. Stefan Pöhlmann. For this study, the spike sequence was cloned into pcDNA3.1(+) vector. Plasmid encoding for SARS-CoV-2 Omicron BA.1 spike was kindly provided by Prof. Dr. Ralf Bartenschlager and purchased from Addgene (#180375). Empty pcDNA3.1(+) vector was used as a negative control. Immunofluorescence staining was obtained by using rabbit anti spike antibody (Abcam, ab272504) for Wuhan-Hu-1 spike and mouse anti-spike for Omicron BA.1 spike (BIOZOL, GTX632604). Secondary antibodies, Alexa Fluor 488 goat anti-rabbit or anti-mouse (Invitrogen, A11034 and A11029), were used. For plasmid DNA transfection, TransIT-LT1 Transfection Reagent was used according to the manufacturer’s protocol (Mirus Bio LLC).

### Cell viability assay

VeroE6 or A549 ACE2+ cells were seeded into 24-well plates (2.5×10^4^ cells/well) in the culture medium descibed above and treated with heparin, unsulfated parental K5, K5OSH and K5NOSH (from 100 µM to 10 nM) at 37°C for 48 h. Cell viability was estimated by measuring the ATP levels using CellTiter-Glo (Promega, Madison, WI, USA). All the experiments were repeated twice.

### Syncytia formation assay and automated detection of syncytia

VeroE6 cells were seeded in a 24-well plate (glass bottom) at a seeding density of 0.04×10^6^ cells/well. Cells were transfected with 500 ng DNA/well. Empty pcDNA3.1(+) vector was used as a negative control. Heparin or K5 derivatives were added after washing the cells 2x with medium 4-6 hours post-transfection (hpt). Cells were washed 2x with PBS at 24 hpt and fixed by 4% formaldehyde (PFA) (Science Services, E15710) diluted in PBS for 15 min at room temperature (RT). Cells were washed 2x with PBS and permeabilized with 0.25% Triton X-100 in PBS for 10 min at RT. Cells were washed 3× with PBS and blocked with 2.5% lipid-free BSA in 0.1% Tween-20 in PBS (PBS-T) for 1 h at RT. After blocking and washing, cells were incubated with primary antibody solutions prepared by diluting the antibody 1:1000 in 1% lipid-free BSA in PBS-T (dilution buffer) for 1 h at RT. Cells were washed 3× with PBS and incubated with secondary antibody solutions prepared by diluting antibodies 1:500 in dilution buffer for 1h at RT in the dark. Cells were washed 3x with PBS and incubated in DAPI (Sigma-Aldrich, D9542, 1:1000 dilution in PBS) for 1 min at RT. After washing 3x with PBS, plates were imaged in an automated fashion using Cell Discoverer 7 (Zeiss, state, city) with 20x (PLAN-Apo, NA 0.7, air) objective and Zeiss Axiocam 712 mono camera. The entire surface of each well was imaged by tiled acquisition in 3 channels (385 nm LED excitation coupled with 425/30 emission filter for DAPI detection, 470 nm LED excitation coupled with 513/30 emission filter for the spike protein detection and a bright field channel). Individual, tiled images were subsequently stitched covering the entire surface of the well containing approx. 40,000 cells per well on average. To measure the extent of syncytia formation in an unbiased way, we employed an automated image analysis pipeline using the Arivis Vision 4D software (Zeiss). First, we segmented nuclei by applying the Cellpose^51^, a pre-trained machine learning model “CP” on the DAPI channel, which resulted in successful nuclei detection even in situations where nuclei were very close to each other, as it is often the case within the syncytia. To detect the boundaries of cells expressing different spike mutants, we trained a dedicated semantic deep-learning model using the Arivis Cloud (Zeiss) platform. For training, we used 10 randomly selected 1000×1000 px regions of interest from one well and all 3 acquired channels. Two different models were trained for the detection of the Wuhan-Hu-1 and the Omicron BA.1 spike variants. Segmented objects in close proximity were separated by the application of the watershed algorithm as implemented in the “Splitting” module of the Arivis Vision 4D software. Only those objects larger than 420 µm^2^ were considered. Syncytia was defined as any cellular object larger than 2,000 μm^2^ and containing at least 2 nuclei. In the final step, all nuclei of the spike-expressing cells were classified as either “in” or “out” of syncytia using the “syncytia criterion” mentioned above. From this analysis, “fusogenicity” was calculated as “% of nuclei in syncytia” by dividing the number of nuclei in syncytia by the total number of nuclei in spike-expressing cells. The same automated acquisition and analysis pipeline was applied to all the experiments, ensuring an unbiased approach and comparable results.

### Infection assay

#### Virus

Infections were carried out as previously described ^52, 53^ using the clinical SARS-CoV-2 isolates belonging to B.1 (GISAID accession number: EPI_ISL_1379197) or Omicron BA.1 (GISAID accession number: EPI_ISL_15700833) lineages ^54^. The viruses were propagated in Vero E6 cells and the viral titer was determined by a standard plaque assay. All the experiments were performed with a single viral inoculum. All the infection experiments were carried out in a biosafety level-3 (BLS-3) laboratory at a Multiplicity of Infection (MOI) of 0.01.

#### Evaluation of antiviral efficacy

VeroE6 or A549 ACE2+ cells were grown to 80%-90% confluence and infected at 37°C for 1 h with the SARS-CoV-2 isolates at a MOI of 0.01. Then, the virus was removed and cells were washed with PBS at 37°C and cultured with media containing 2% FBS in the presence or the absence of heparin, unsulfated parental K5, K5OSH and K5NOSH (from 10 µM to 10 nM). Supernatants were collected for further analysis 48 h post-infection (p.i). Mock-infected cells were processed exactly as the SARS-CoV-2-infected ones, except they were not exposed to the virus.

#### Plaque Assay

Vero E6 cells were seeded at a density of 5×10^5^ cells/well in a 12-well plate and incubated at 37°C for 24 h. Supernatants from infected cells were serially diluted in DMEM without FBS and added to the cells. After 1 h incubation, media were removed and cells were washed with PBS at 37°C. Then cells were covered with an overlay consisting of DMEM with 0.4% SeaPlaque (Lonza, Basel, Switzerland). Cells were further incubated at 37°C for 48 h and then fixed with 10% formaldehyde at room temperature for 3 h. Formaldehyde and agarose overlay were removed. Cells were then stained with crystal violet (1% w/v in a 20% ethanol solution), and the viral titer (Plaque Forming Unit, PFU/mL) of SARS-CoV-2 was determined by counting the number of plaques. All the experiments were repeated twice.

### Data analysis

GraphPad Prism version 8.4 (GraphPad Software, Boston, Massachusetts USA, www.graphpad.com) was used to perform non-linear regression analysis, plot the dose-response curves and calculate the half-maximal inhibitory concentration (IC_50_) values for each compound. Student’s t-test or one-way ANOVA analysis of variance was performed using Microsoft Excel. P value was determined by Mann-Whitney test (* P < 0.05; ** P< 0.005. *** P < 0.0005).

## Data availability statement

All data that support the findings of this study are publicly available in the Zenodo repository. BiorXiv preprint server XXX.

## Acknowledgments

Glycores 2000. S.r.l for providing the heparin and K5 derivatives.

## Author Contributions

M.M. methodology, validation, SPR analysis. C.U. furin cleavage assays. L.Z.: methodology, validation, syncytia assays, writing, review & editing. P.O. resources (glycosaminoglycans). A.Z. infection assays. A.C. funding acquisition. F.C. infection assays, writing original draft. V.L.: methodology, validation. P.C. methodology, validation, review & editing, supervision, funding acquisition; R.C.W.: conceptualization, resources, review & editing, supervision, funding acquisition. M.R.: conceptualization, resources, writing, review & editing, funding acquisition. G.P.: conceptualization, methodology, validation, investigation, simulations, writing original draft, visualization, review & editing.

## Supporting information

### Supporting Figures

**Fig. S1.**
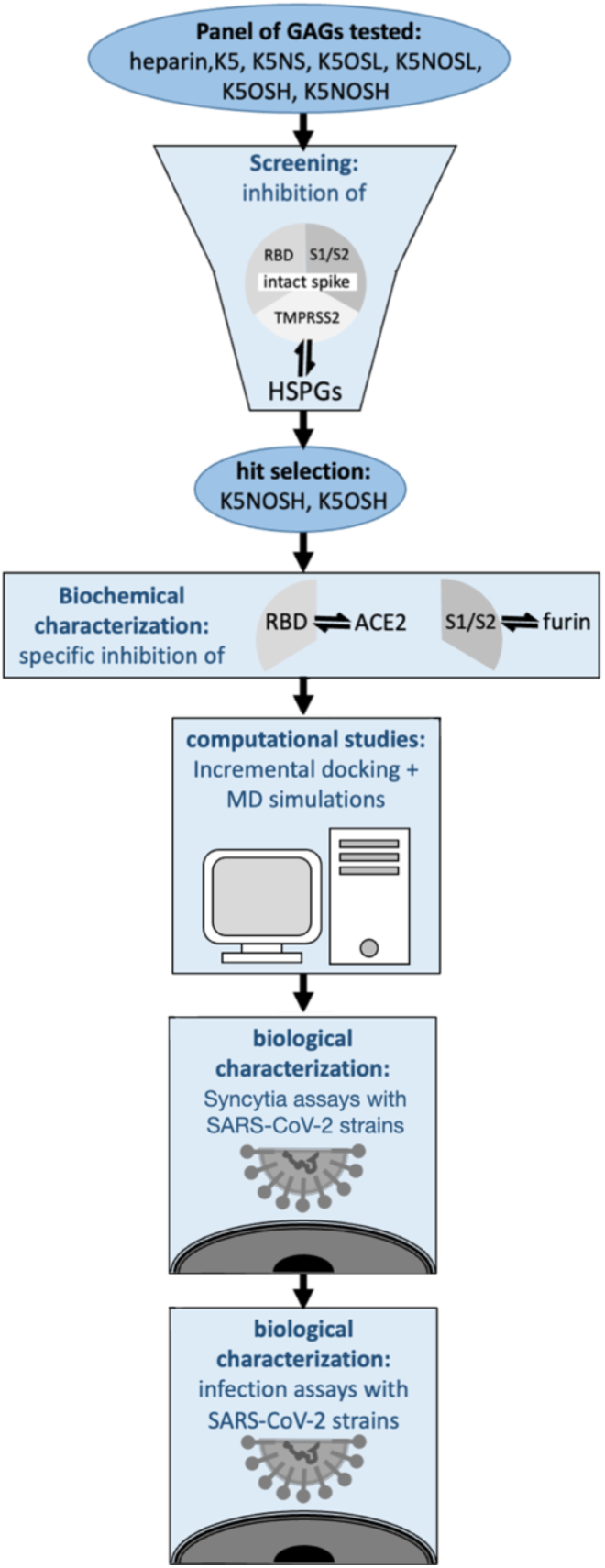
Workflow of the multidisciplinary approach adopted for the identification of anti-SARS-CoV-2 sulfated K5 derivatives and for the characterization of their mechanisms of action.

**Fig. S2.**
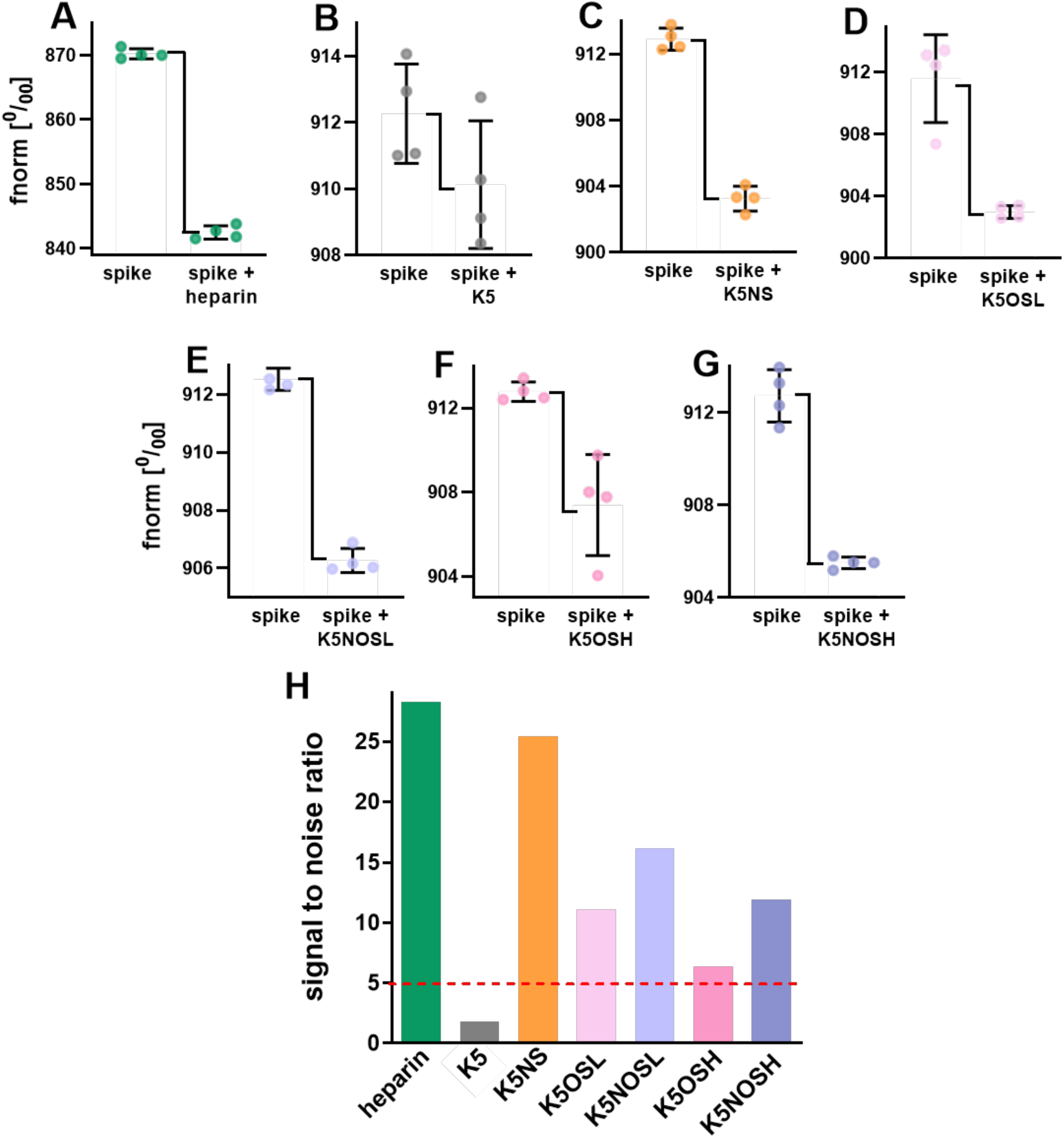
Results of Microscale Thermophoresis (MST) binding check assay. Binding check results for the interaction of spike (20 nM) and the indicated GAGs (15 µM) in 50 mM Tris buffer pH 7.4, supplemented with 150 mM NaCl, 10 mM MgCl_2_ and 0.05% Tween-20. An MST on-time of 1.5 sec was used for analysis. (n = 4. **A-G**: Fnorm values of spike alone and in complex with the indicated GAGs ADP (complex). **H**: Signal-to-noise ratio obtained from the normalized fluorescence (Fnorm ^0^/_00_) evaluation shown above. The horizontal red line indicates the signal-to-noise ratio value designated as the cut-off between binding and non-binding molecules following the manufacturer’s instructions.

**Fig. S3.**
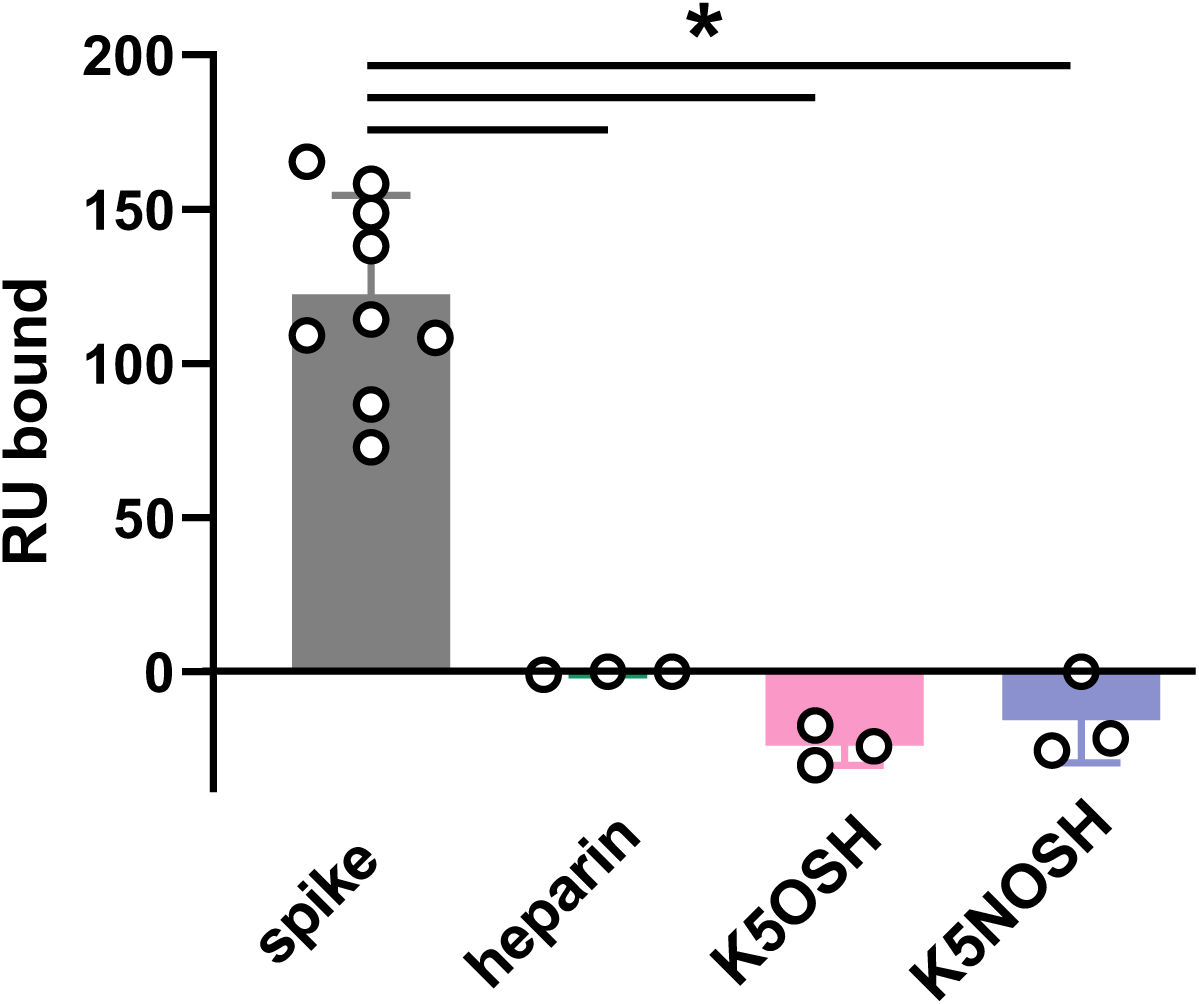
Lack of binding of heparin, K5OSH and K5NOSH to sensorchip-immobilized ACE2. Spike RBD (20 nM) or the indicated GAGs (100 μM) were injected over the ACE2 containing biosensor to evaluate their capacity to bind directly to the receptor. * P < 0.05.

**Fig. S4.**
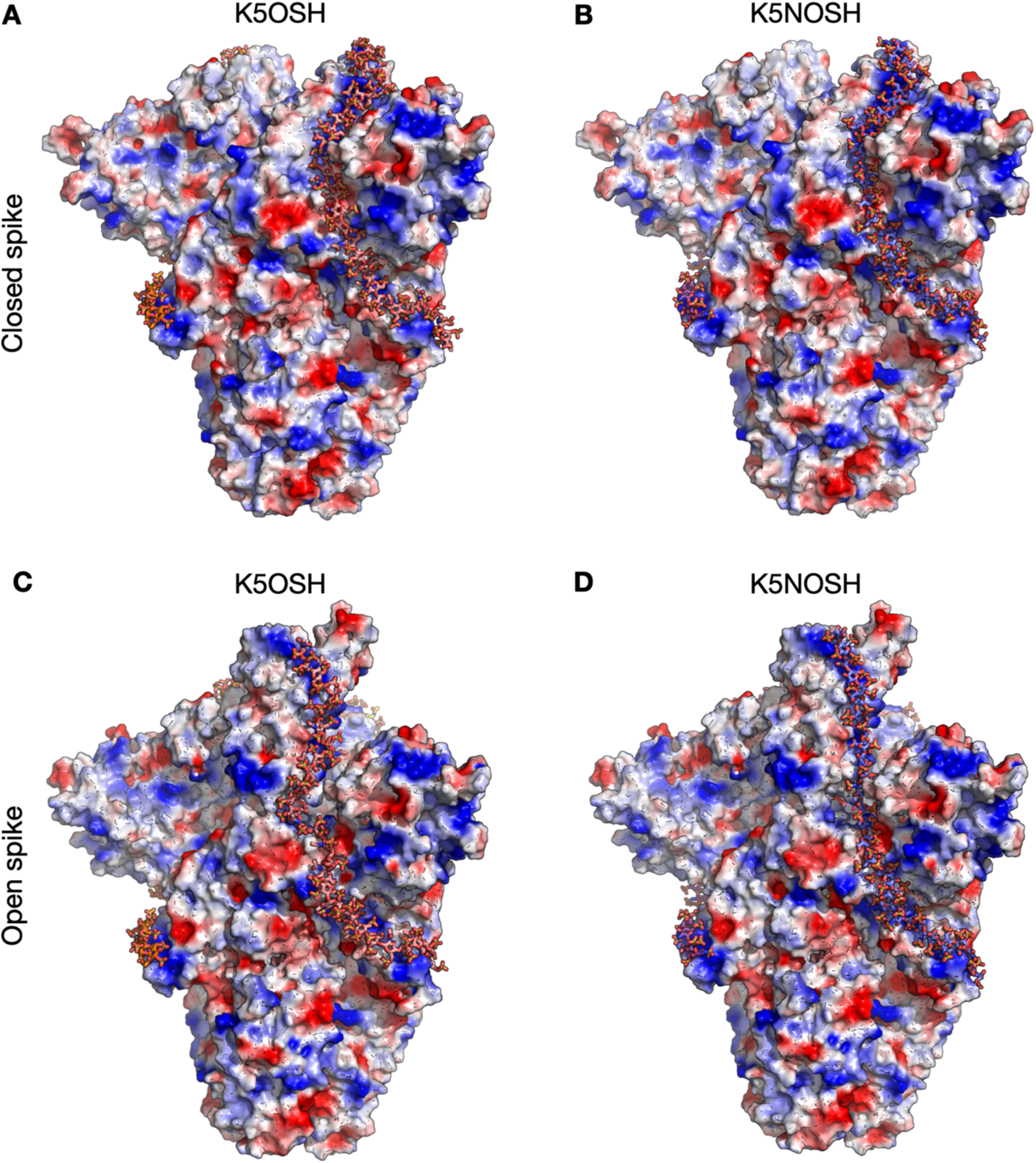
Models of K5 compounds bound to closed and open conformations of the spike glycoprotein. Representative structures of (A-B) closed spike glycoprotein with 3 polysaccharide chains of K5OSH (left) or K5NOSH (right) bound, and (C-D) open spike glycoprotein with 3 polysaccharide chains of K5OSH (left) or K5NOSH (right) bound, obtained using the incremental sliding window docking method ^1^. Closed and open spike glycoproteins are shown as molecular surfaces colored according to electrostatic potential, with glycans omitted for clarity. The 31mer K5 chains follow basic paths along the spike surface that span from the RBD at the top of one spike subunit down to the S1/S2 site of an adjacent subunit. They are shown in stick representation colored by element with pink and purple carbons for K5OSH and K5NOSH, respectively.

**Fig. S5.**
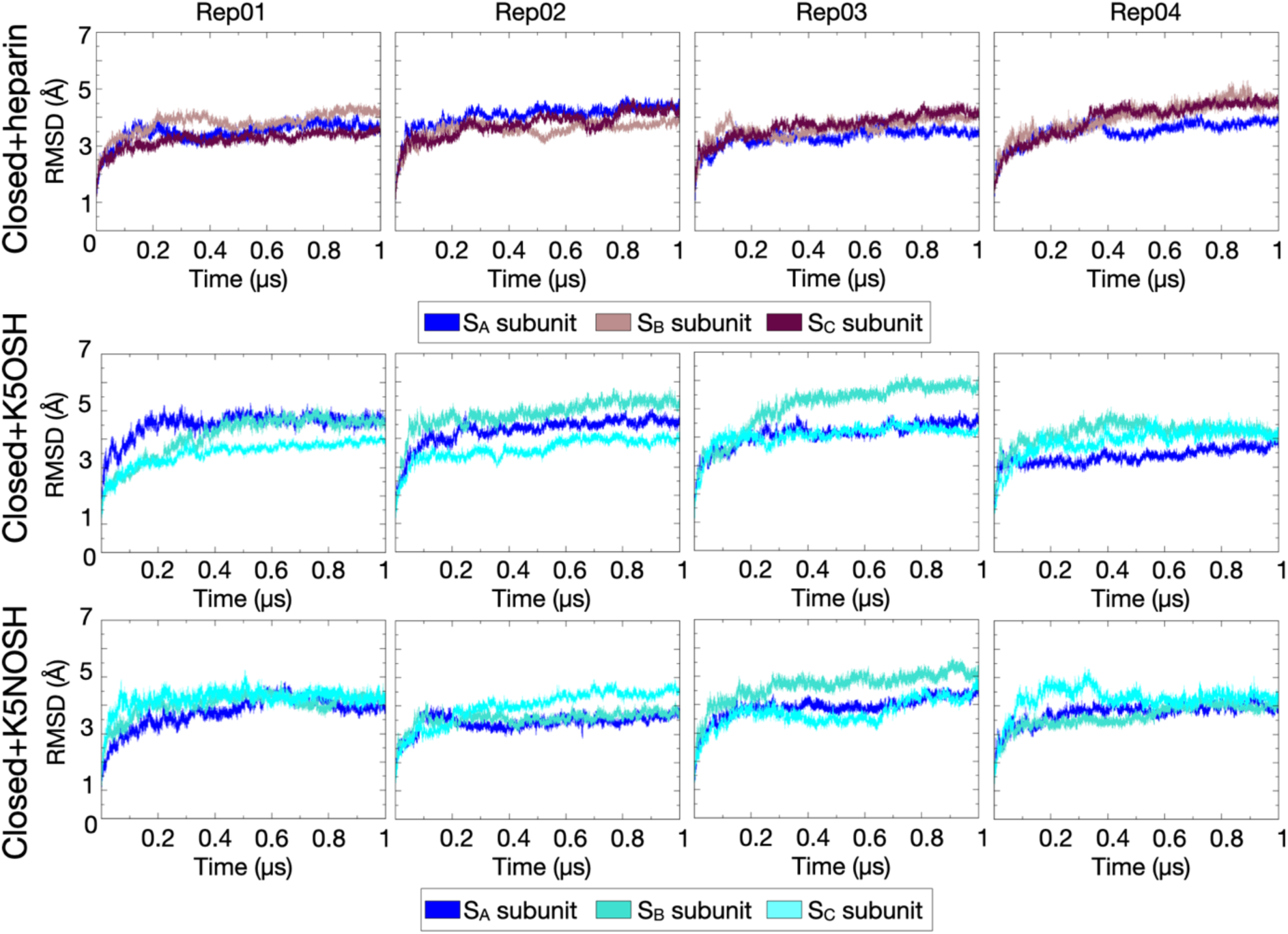
Structural convergence of the closed conformation of spike during the four replica simulations of each complex. Root mean squared deviation (RMSD (Å)) *versus* time (μs) for the systems of closed spike with heparin bound were retrieved from Paiardi et al. ^2^. The four replica MD trajectories of the two simulated systems of closed spike glycoprotein upon K5OSH and K5NOSH engagement are from the current study. The RMSD values for the individual spike subunits - S_A_, S_B_, S_C_ - were calculated for the C-alpha atoms of residues 51-1063 of each spike subunit and are shown in blue, pink, and brown for the spike-heparin complex, and in blue, teal, and cyan for the spike-K5 complexes.

**Fig. S6.**
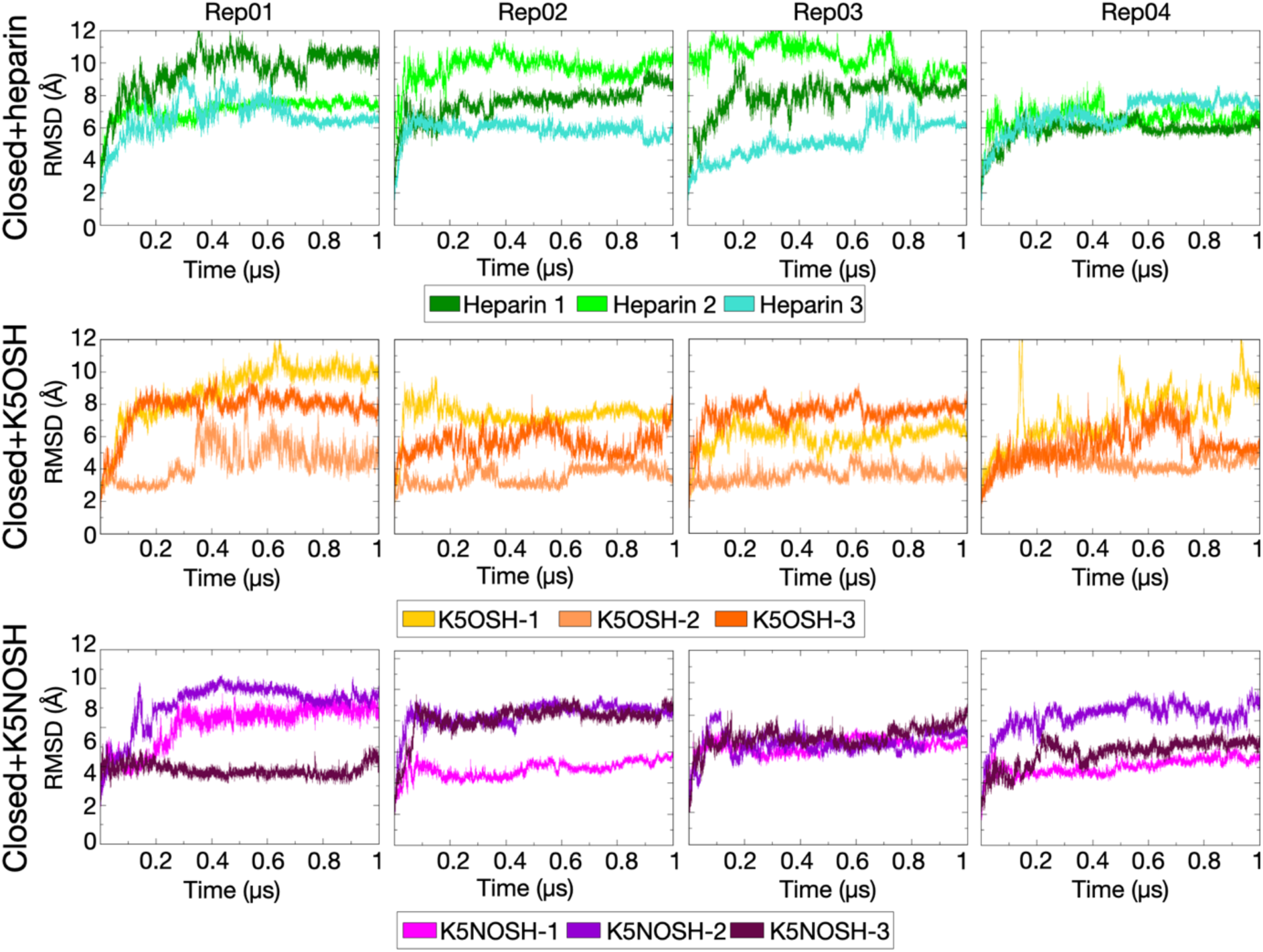
Structural convergence of heparin and the K5 compounds bound to closed spike for the simulated complexes. Root mean squared deviation (RMSD (Å)) *versus* time (μs) of the heparin chains was retrieved from Paiardi et al. ^2^. The four replica MD trajectories of the two simulated systems of closed spike glycoprotein upon K5OSH and K5NOSH engagement are from the current study. RMSD values of all the polyanionic chains were calculated for all the C, N and O atoms in all monosaccharides and are shown in forest, green, and teal for heparin; yellow, orange, and dark orange for K5OSH; magenta, purple, and crimson for K5NOSH.

**Fig. S7.**
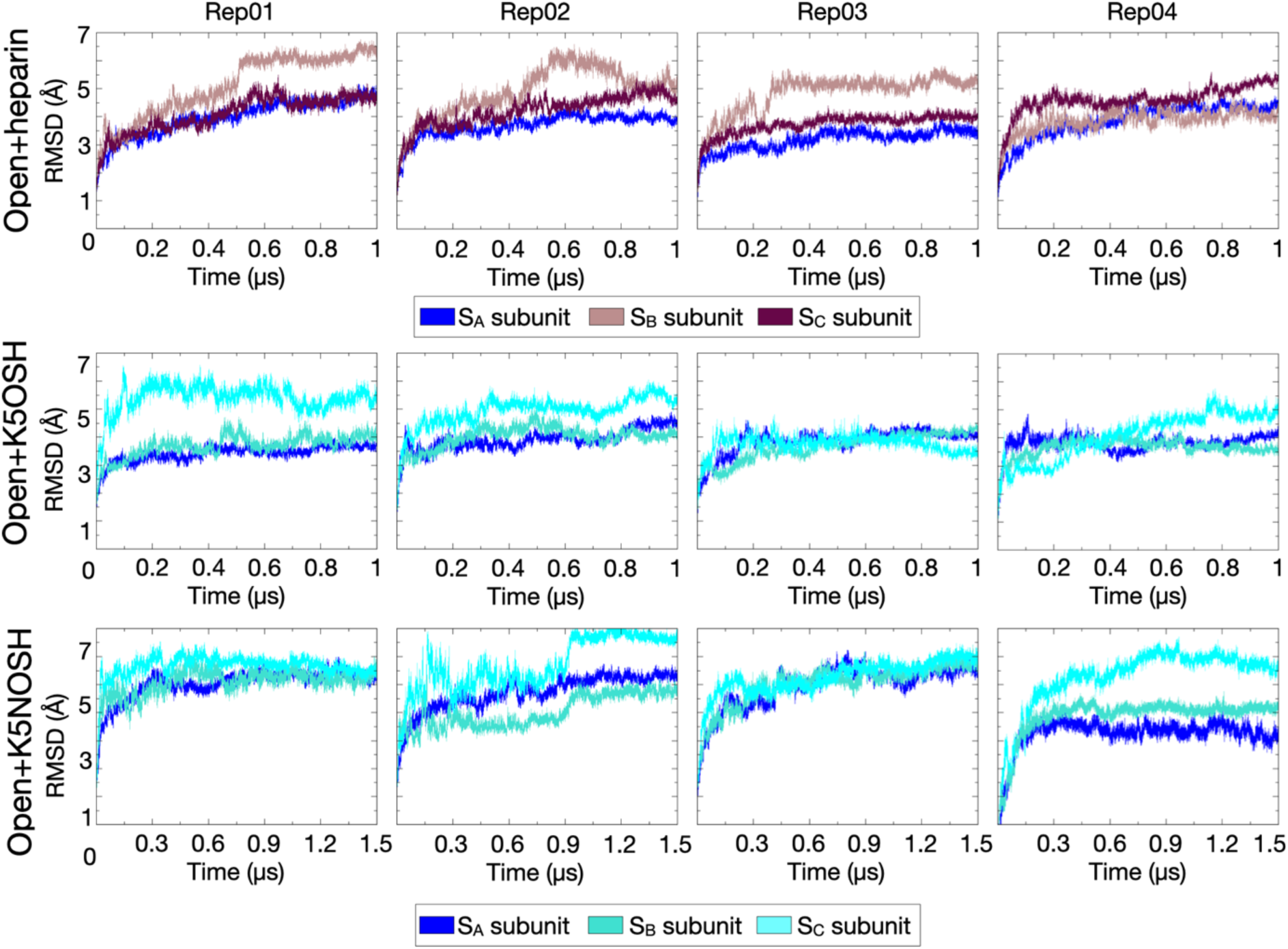
Structural convergence of the open spike during the four replica simulations of each complex. Root mean squared deviation (RMSD (Å)) *versus* time (μs) for the simulated systems of open spike with heparin bound were retrieved from Paiardi et al. ^2^. The four replicas MD trajectories of the two simulated systems of open spike glycoprotein systems upon K5OSH and K5NOSH engagement were from the current study. The RMSD values for the individual subunits - S_A_, S_B_, S_C_ - were calculated for the C-alpha atoms of residues 51-1063 for each spike subunit and are shown in blue, pink, and brown for the spike-heparin complex, and in blue, teal, and cyan for the spike-K5 complexes.

**Fig. S8.**
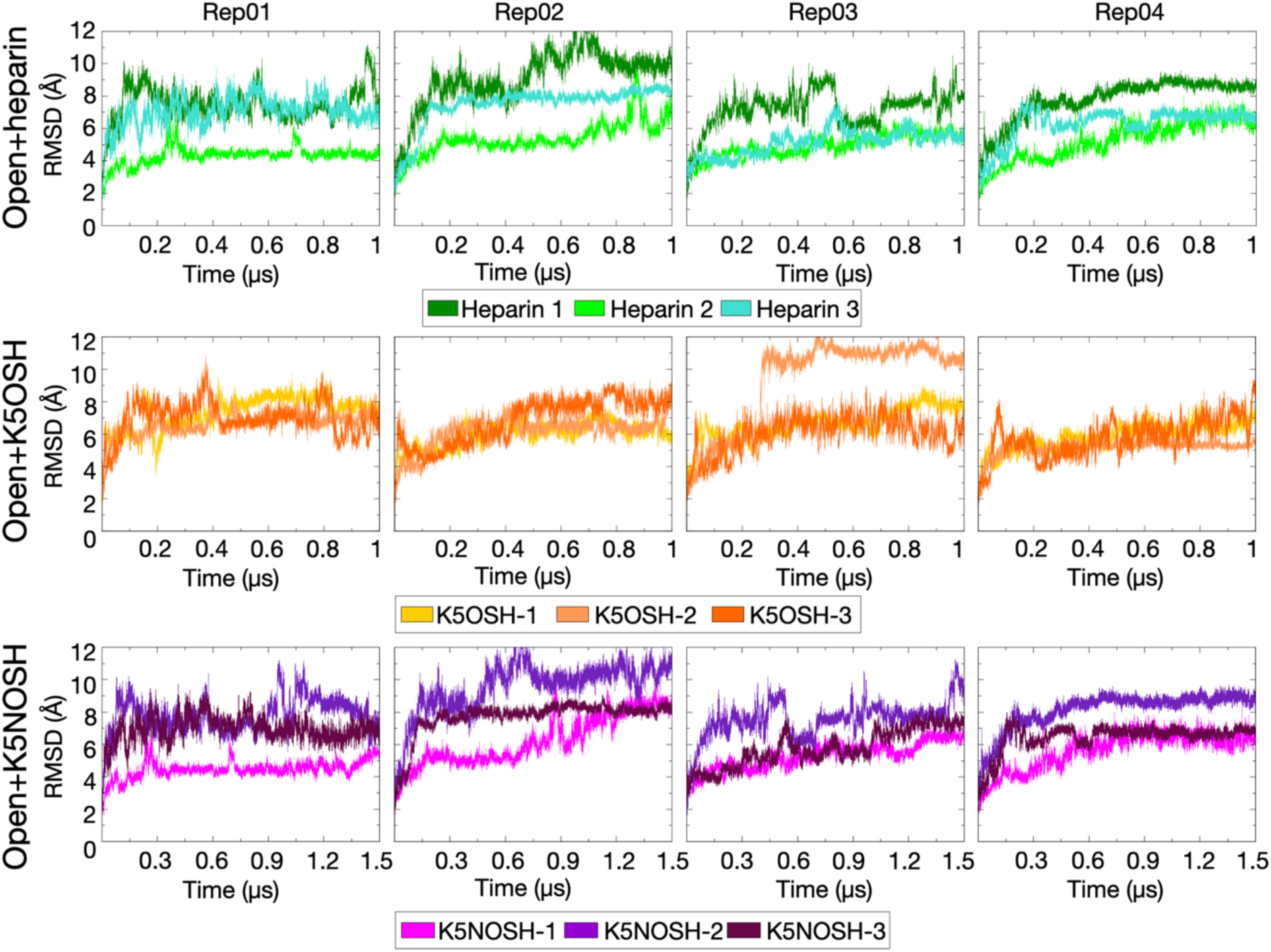
Structural convergence of heparin and the K5 compounds bound to open spike for the simulated complexes. Root mean squared deviation (RMSD (Å)) *versus* time (μs) of heparin chains was retrieved from Paiardi et al. ^2^. The four replicas MD trajectories of the two simulated systems of open spike glycoprotein upon K5OSH and K5NOSH engagement are from the current study. RMSD values of all the polyanionic chains were calculated for all the C, N, and O atoms in all monosaccharides and are shown in forest, green, and teal for heparin; yellow, orange, and dark orange for K5OSH; magenta, purple, and crimson for K5NOSH.

**Fig. S9.**
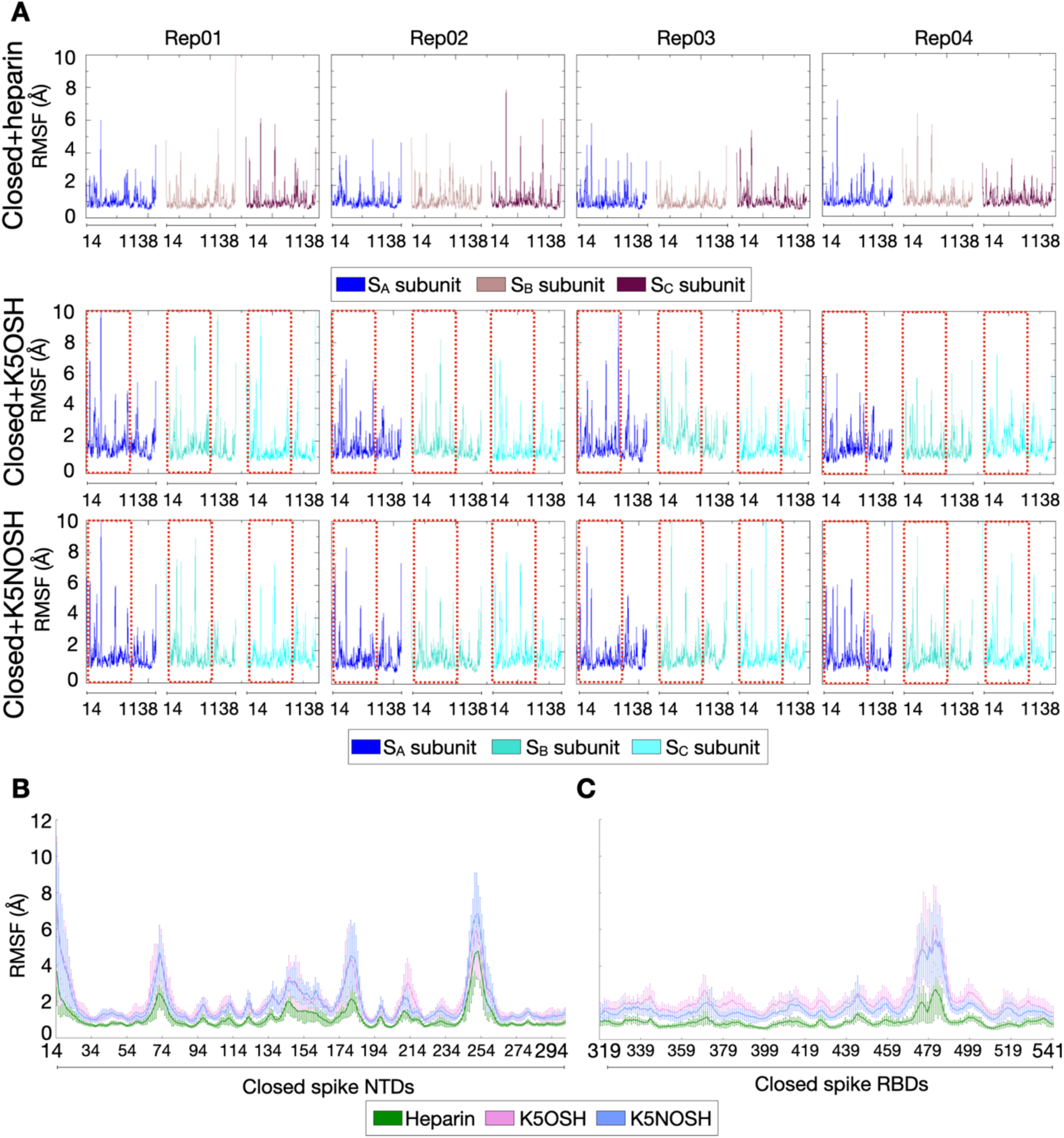
Conformational fluctuation of closed spike in the simulated systems. **(A)**The root mean squared fluctuation (RMSF) (Å) *versus* residue number of the three subunits of the spike upon heparin binding was retrieved from Paiardi et al. ^2^. The four replica MD trajectories of the two simulated systems in the presence of K5OSH and K5NOSH are from the current study. The RMSF values for the individual subunits - S_A_, S_B_, S_C_, - were calculated for the C-alpha atoms of residues 14-1138 and are shown in blue, pink, and brown for the heparin-spike complexes, and in blue, teal, and cyan for the K5-spike complexes. The red boxes enclose the NTD (residues 14-294) and RBD (residues 319-541) of the closed spike subunits upon K5OSH engagement. The average and standard deviation of the RMSF values for the NTD **(B)** and RBD **(C)** over the replica MD trajectories of the simulated systems upon heparin (green), K5OSH (pink), and K5NOSH (purple) binding are plotted per residue along the sequence.

**Fig. S10.**
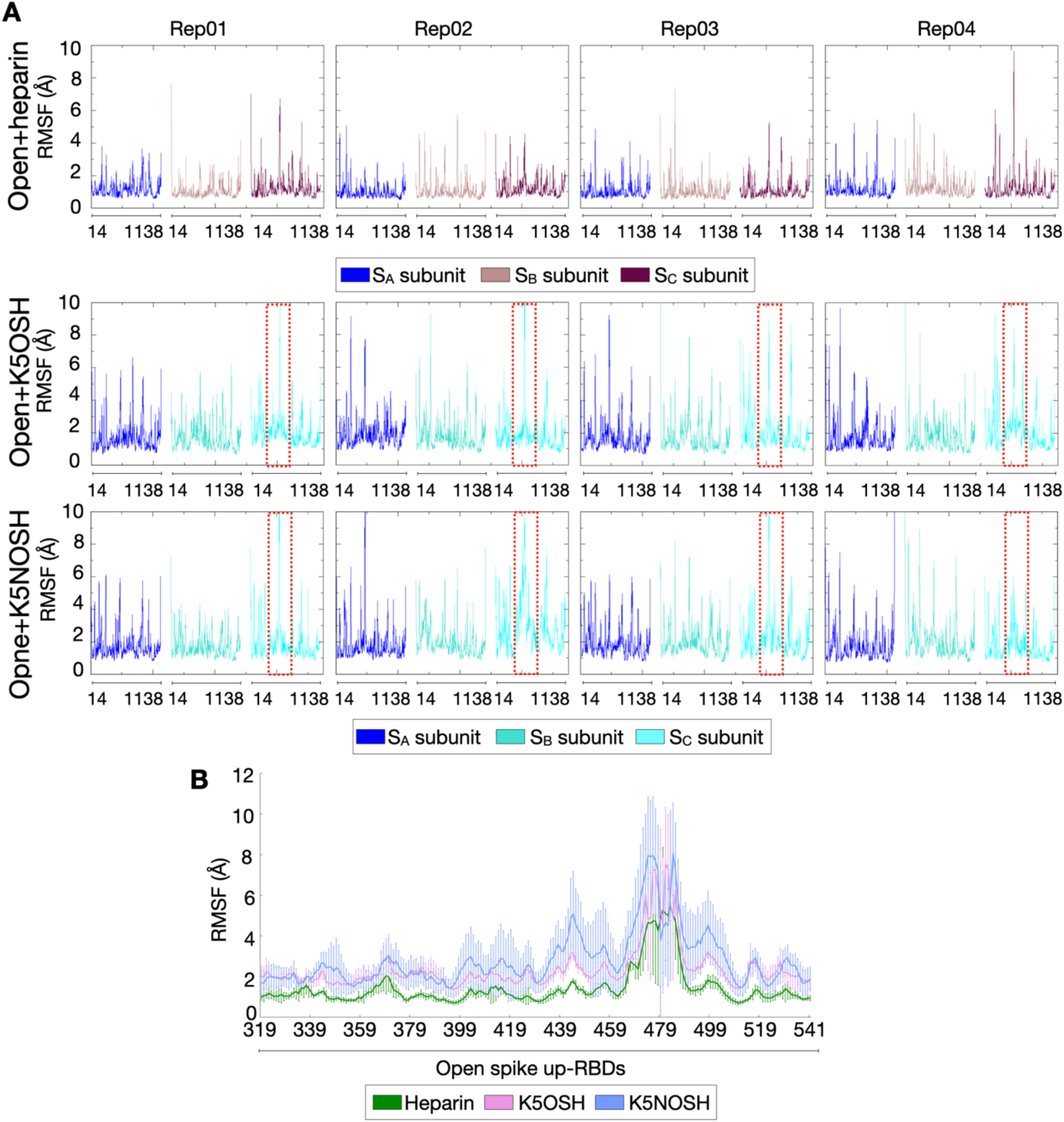
Conformational fluctuation of open spike in the simulated systems. **(A)** The root mean squared fluctuation (RMSF) (Å) *versus* residue number of the three subunits of the spike upon heparin binding was retrieved from Paiardi et al. ^2^. The four replica MD trajectories of the two simulated systems in the presence of K5OSH and K5NOSH are from the current study. The RMSF values for the individual subunits - S_A_, S_B_, S_C_ - were calculated for the C-alpha atoms of residues 14-1138 and are shown in blue, pink, and brown for the heparin-spike complexes, and in blue, teal, and cyan for the K5-spike complexes. The red boxes enclose the RBD (residues 319-541) of the open spike S_C_ (up-RBD) subunits upon K5NOSH engagement. The average and standard deviation of the RMSF values for the spike S_C_ RBD **(B)** over the replica MD trajectories of the simulated systems upon heparin (green), K5OSH (pink), and K5NOSH (purple) binding are plotted per residue along the sequence.

**Fig. S11.**
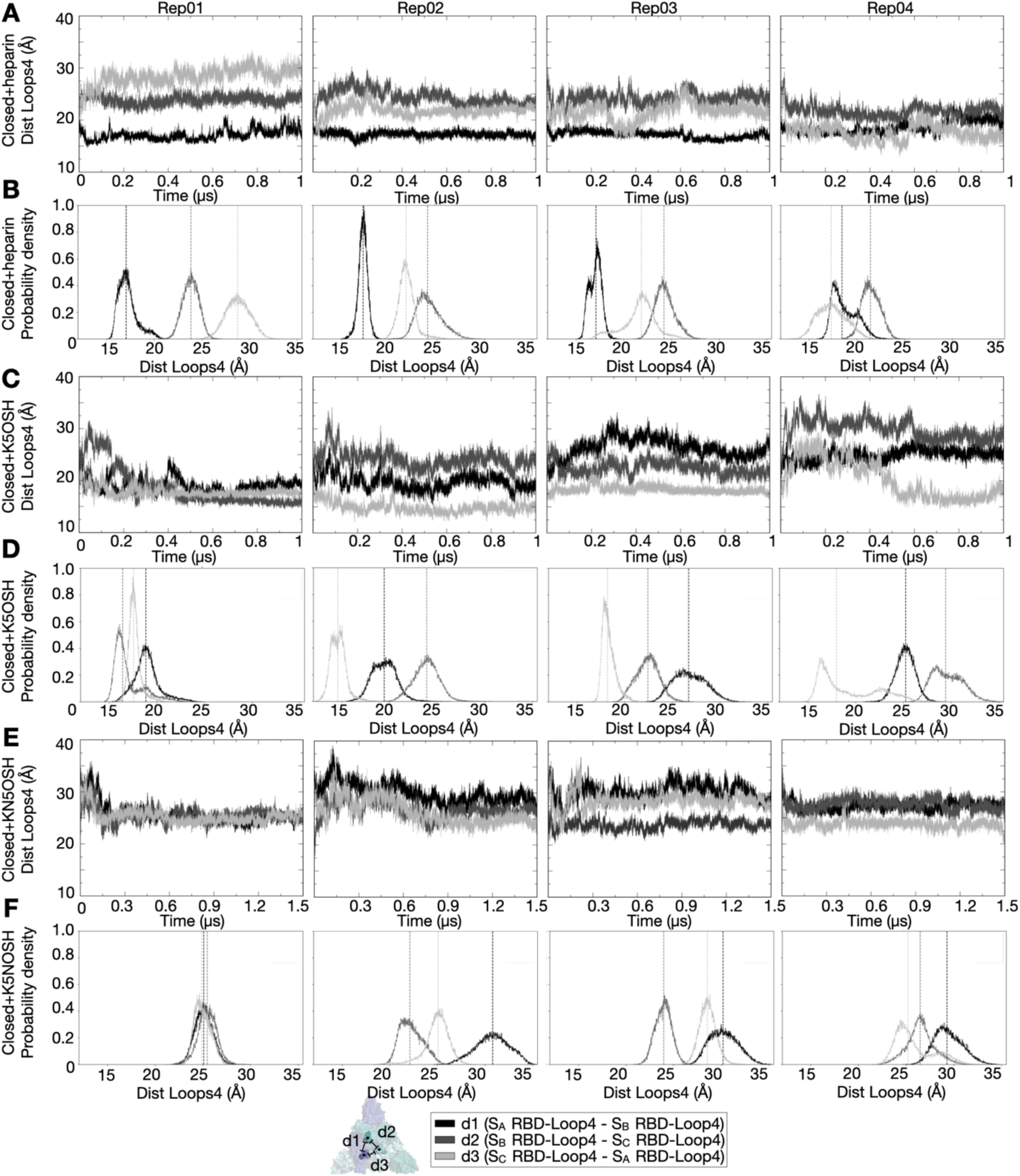
Packing of the closed spike upon heparin and K5 engagement. **(A-C-E)** The distance between two centroids *versus* time (μs) and **(B-D-F)** the probability density *versus* the distance between centroids (Å) for the simulated systems of closed spike with **(A-B)** heparin bound for trajectories from Paiardi et al.^2^ and for MD trajectories simulated in the current study for the two closed spike glycoprotein systems upon **(C-D)** K5OSH and **(E-F)** K5NOSH engagement. Centroids for the individual subunits - S_A_, S_B_, S_C_ - were calculated for residues of RBD Loop4 (residues 495-516). **(A-C-E)** The distance between two centroids computed along the simulation and **(B-D-F)** the probability density computed on the converged MD trajectories (from 0.2 to 1 μs) are shown in black, dark grey and light grey for the distances: S_A_ RBD Loop4 - S_B_ RBD Loop4 (d1), S_B_ RBD Loop4 – S_C_ RBD Loop4 (d2) and S_C_ RBD Loop4 – S_A_ RBD Loop4 (d3), respectively. **(B-D-F)** The dashed lines showing the mean values for the single distribution are colored according to the respective distance (black, dark grey and light grey).

**Fig. S12.**
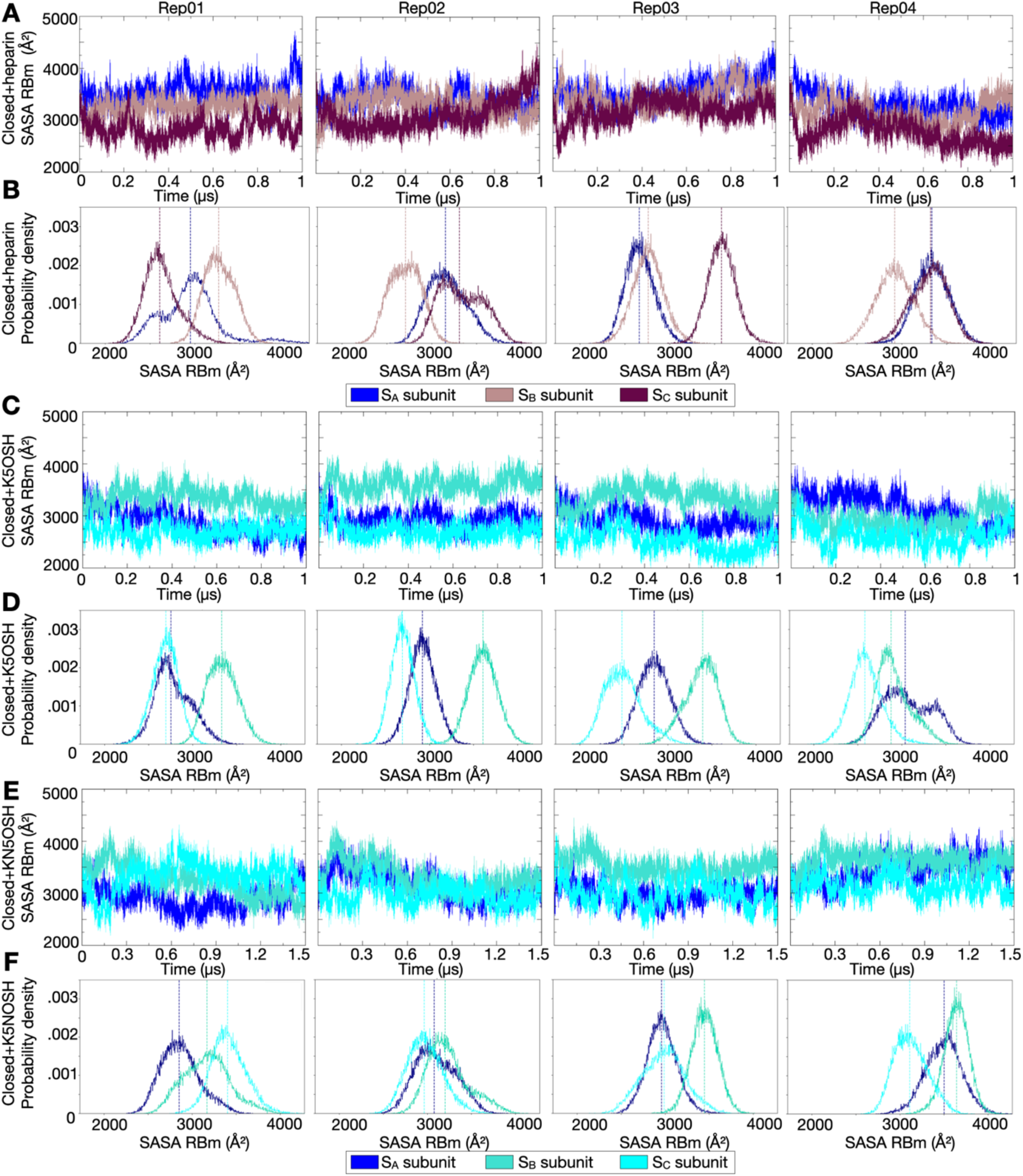
Exposure of the receptor binding motif (RBm) during the MD simulations. **(A-C-E)** Solvent Accessible Surface Area (SASA) (Å^2^) of the RBm versus time (µs) and **(B-D-F)** the probability density *versus* SASA of RBm (Å^2^) for the simulated systems of **(A-B)** closed spike with heparin bound (from Paiardi et al.^2^) and for the four replica MD trajectories of the two systems of closed spike glycoprotein with **(C-D)** K5OSH and **(E-F)** K5NOSH bound simulated in the current study. **(A-C-E)** The SASA is computed along the simulations while **(B-D-F)** the probability density is computed on the converged MD trajectories (from 0.2 to 1 μs) and is shown for each subunit - S_A_, S_B_, S_C_ - in blue, pink, and brown for systems with heparin bound, in blue, teal, and cyan for systems with K5 compounds bound. **(B-D-F)** The dashed lines showing the mean values for the single distribution are colored according to the SASA of the respective subunit.

**Fig. S13.**
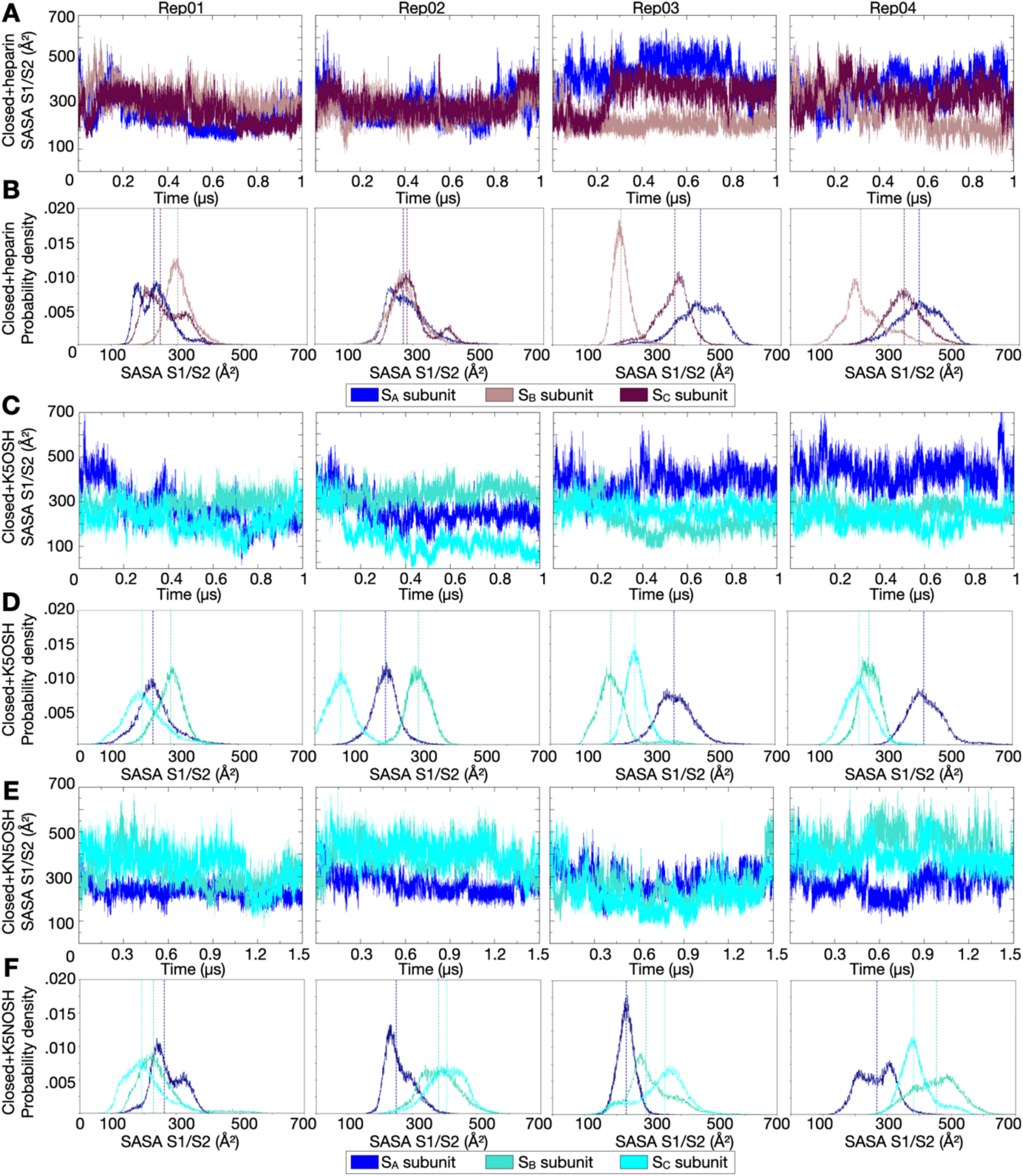
Exposure of the S1/S2 multifunctional site during the MD simulations. **(A-C-E)** Solvent Accessible Surface Area (SASA) (Å^2^) of the S1/S2 site versus time (µs) and **(B-D-F)** the probability density *versus* SASA of the S1/S2 site (Å^2^) for the simulated systems of **(A-B)** closed spike with heparin bound (from Paiardi et al.^2^) and for the four replica MD trajectories of the two systems of closed spike glycoprotein with **(C-D)** K5OSH and **(E-F)** K5NOSH bound simulated in the current study. **(A-C-E)** The SASA is computed along the simulations while **(B-D-F)** the probability density is computed on the converged MD trajectories (from 0.2 to 1 μs) and is shown for each subunit - S_A_, S_B_, S_C_ - in blue, pink, and brown for systems with heparin bound, in blue, teal, and cyan for systems with K5 compounds bound. **(B-D-F)** The dashed lines showing the mean values for the single distribution are colored according to the SASA of the respective subunit.

**Fig. S14.**
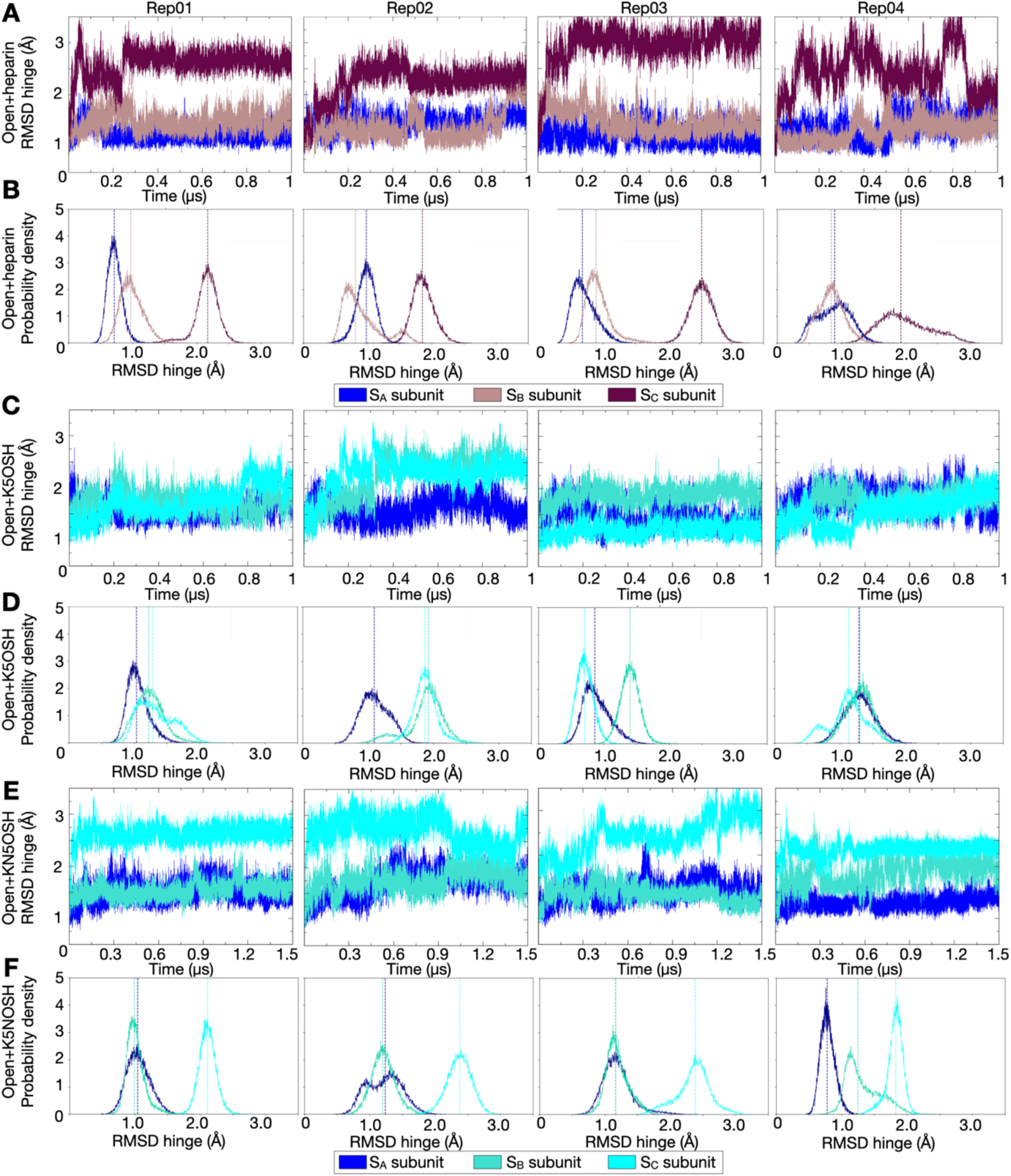
Conformational change of the hinge region of the up-RBD during the simulations. **(A-C-E)** The root mean square deviation (RMSD) (Å) of the hinge region versus time (µs) and **(B-D-F)** the probability density *versus* RMSD of the hinge region (Å^2^) for the simulated systems of **(A-B)** closed spike with heparin bound (from Paiardi et al.^2^) and for the four replica MD trajectories of the two systems of closed spike glycoprotein with **(C-D)** K5OSH and **(E-F)** K5NOSH bound simulated in the current study. **(A-C-E)** The RMSD values were calculated for the C-alpha atoms of residues 527-529 a long the simulations while **(B-D-F)** the probability density was computed on the converged MD trajectories (from 0.2 to 1 μs) and is shown for each subunit - S_A_, S_B_, S_C_ - in blue, pink, and brown for systems with heparin bound, in blue, teal, and cyan for systems with K5 compounds bound. **(B-D-F)** The dashed lines showing the mean values for the single distribution are colored according to the SASA of the respective subunit.

**Fig. S15.**
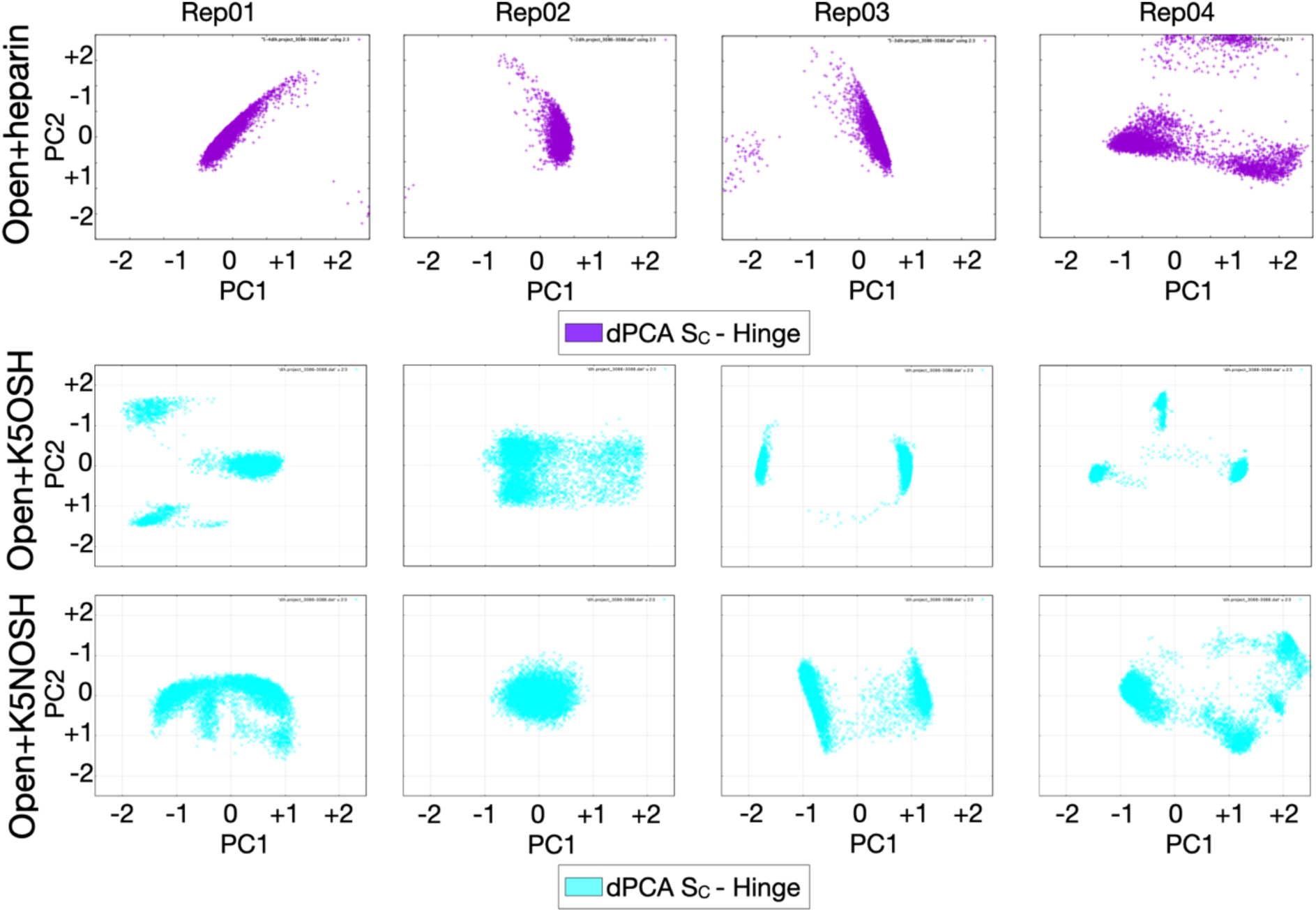
Rearrangement of the RBD-hinge upon K5 engagement. Results of dihedral Principal Component Analysis (dPCA) calculated for the Phi/Psi angles of the RBD-hinge region (residues 527-529) of the S_C_ subunit for the simulated systems of spike with heparin bound from Paiardi et al.^2^ are colored magenta and those for the four replica MD trajectories of the two open spike glycoprotein systems upon K5OSH and K5NOSH binding are colored cyan. Principal components (PC) 1 and 2 are plotted on the x and y axes, respectively.

**Fig. S16.**
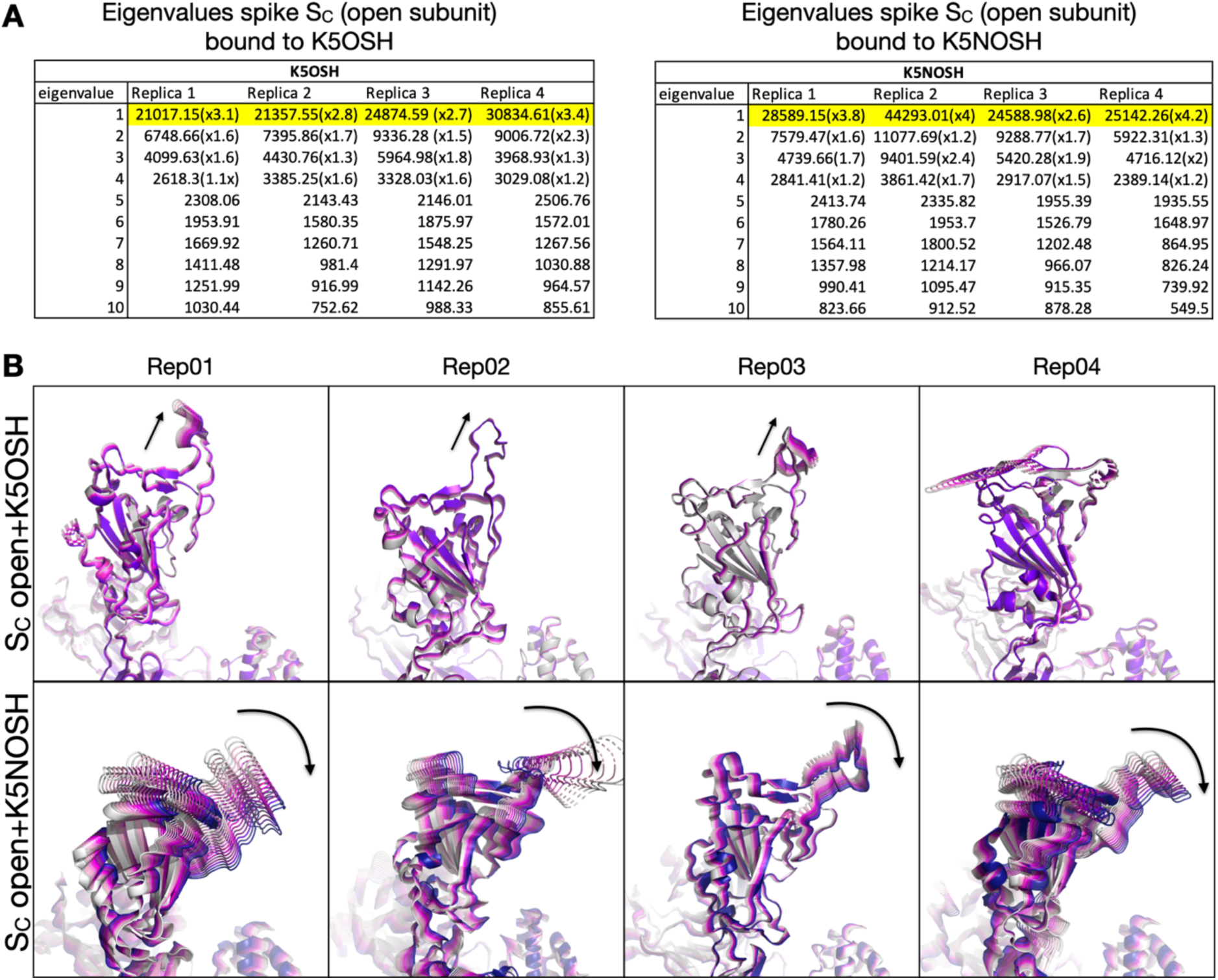
Differences in the closing motion of the open spike RBD in the presence of K5 compounds. Essential Dynamics (ED) analysis on the S_C_ subunit (up-RBD) for the two replica MD trajectories of open spike bound to K5OSH or K5NOSH polysaccharides. (A) Values of the first ten eigenvalues for the up-subunit of spike bound to K5OSH or K5NOSH polysaccharides in the four replica simulations. The first eigenvector dominates the motion in all cases. Multipliers within the brackets denote the magnitude of each eigenvalue with respect to the next eigenvalue. (B) Plot of the direction of motion of the first eigenvector on the RBD of the S_C_ subunit which can be seen to differ between spike bound to K5OSH and to K5NOSH polysaccharides. The second replica of the system with K5NOSH has a less pronounced closing movement than the other replicas.

**Fig. S17.**
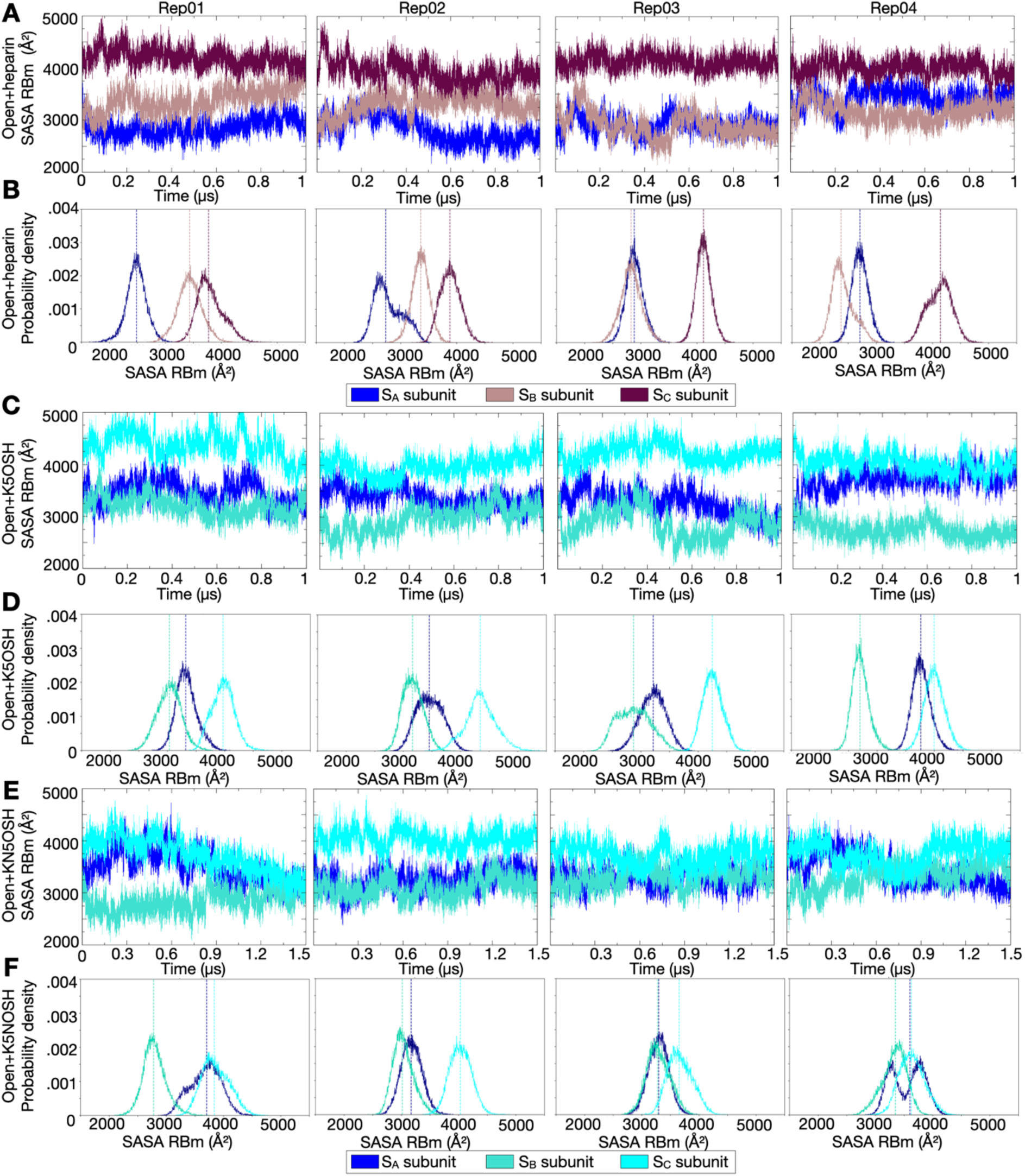
Exposure of the receptor binding motif (RBm) during the MD simulations. **(A-C-E)** Solvent Accessible Surface Area (SASA) (Å^2^) of the RBm versus time (µs) and **(B-D-F)** the probability density *versus* SASA of RBm (Å^2^) for the simulated systems of open spike with heparin bound from Paiardi et al.^2^ and for the four replica MD trajectories of the two systems of open spike glycoprotein with K5OSH and K5NOSH bound simulated in the current study. The SASA is shown for each subunit - S_A_, S_B_, S_C_ - in blue, pink, and brown for heparin-spike complexes and in blue, teal, and cyan for K5-spike complexes. **(A-C-E)** The SASA is computed along the simulations while **(B-D-F)** the probability density is computed on the converged MD trajectories (from 0.2 to 1 μs) and is shown for each subunit - S_A_, S_B_, S_C_ - in blue, pink, and brown for systems with heparin bound, in blue, teal, and cyan for systems with K5 compounds bound. **(B-D-F)** The dashed lines showing the mean values for the single distribution are colored according to the SASA of the respective subunit.

**Fig. S18.**
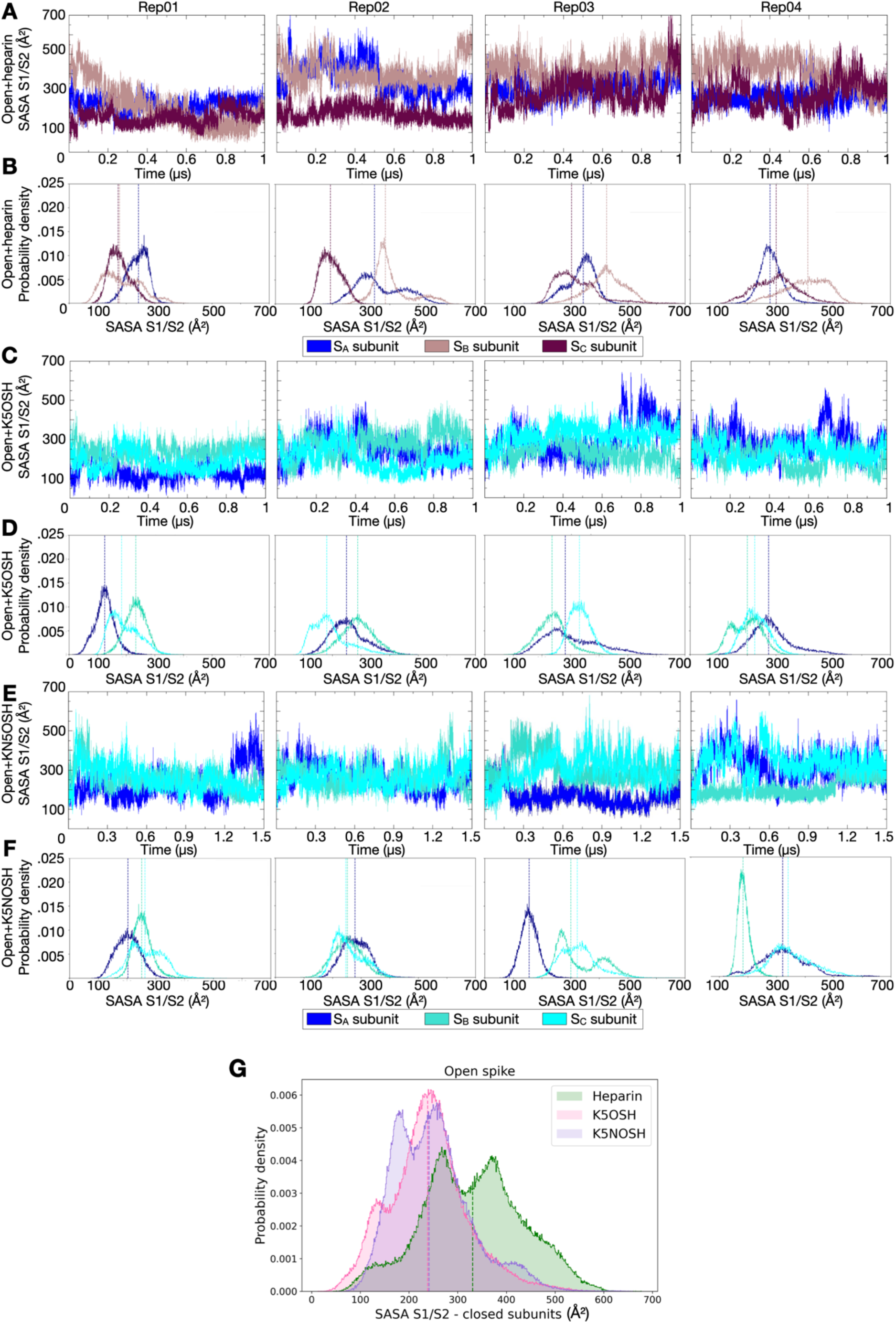
Exposure of the S1/S2 multifunctional site during the MD simulations. **(A-C-E)** Solvent Accessible Surface Area (SASA) (Å^2^) of the S1/S2 site versus time (µs) and **(B-D-F)** the probability density *versus* SASA of the S1/S2 site (Å^2^) for the simulated systems of **(A-B)** open spike with heparin bound (from Paiardi et al.^2^) and for the four replica MD trajectories of the two systems of closed spike glycoprotein with **(C-D)** K5OSH and **(E-F)** K5NOSH bound simulated in the current study. **(A-C-E)** The SASA is computed along the simulations while **(B-D-F)** the probability density is computed on the converged MD trajectories (from 0.2 to 1 μs) and is shown for each subunit - S_A_, S_B_, S_C_ - in blue, pink, and brown for systems with heparin bound, in blue, teal, and cyan for systems with K5 compounds bound. **(G)** The probability distribution of the SASA for the S1/S2 of the closed subunits on the open spike (S_A_ and S_B_) was calculated from the converged MD trajectories (from 0.2 to 1 μs) for all the replicas and colored according to systems.

**Fig. S19.**
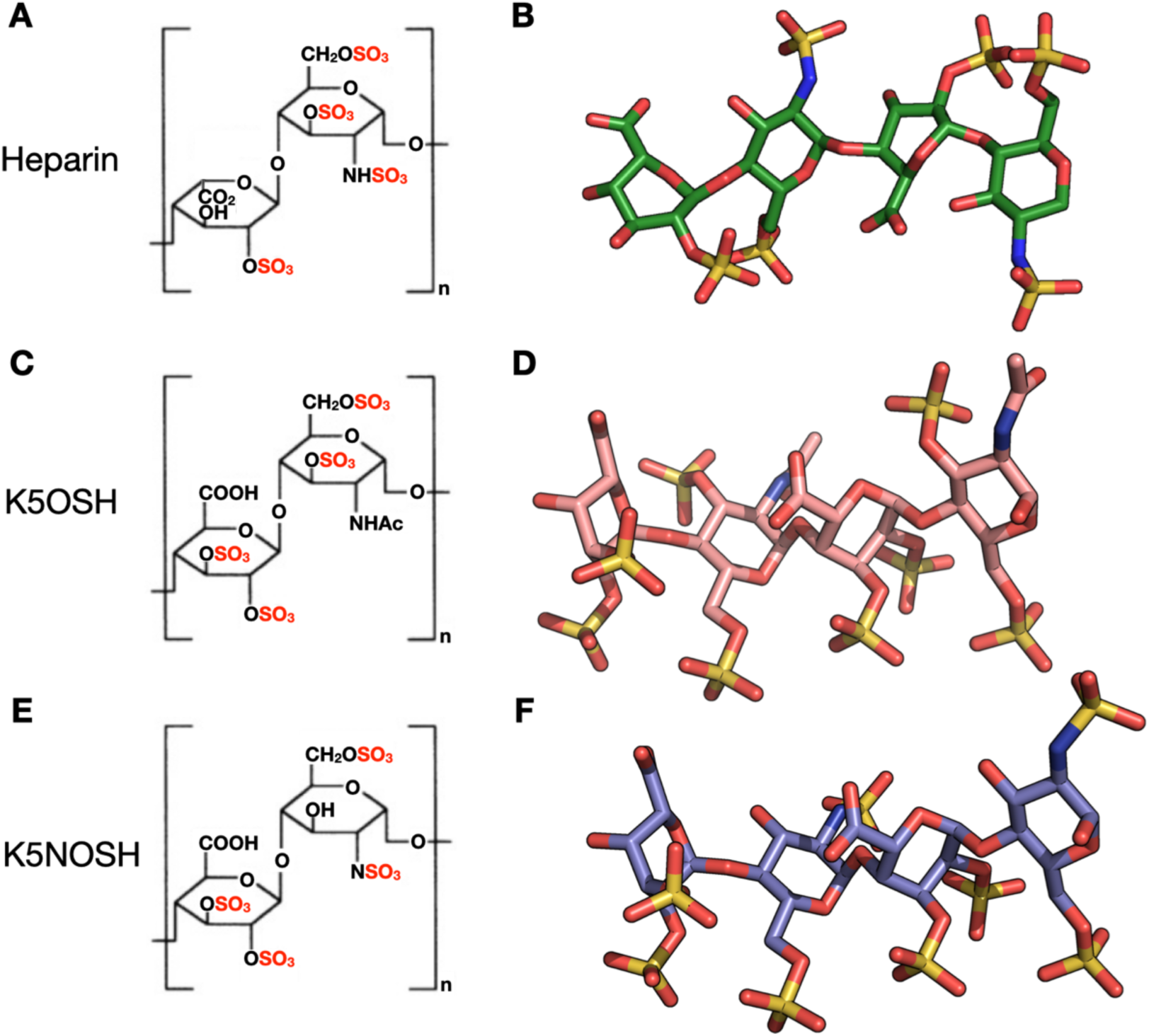
Molecular structures and models of the K5 polysaccharides. 2D representation of the (A) heparin, (C) K5OSH, and (E) K5NOSH disaccharide repeat units used to model the K5NOH and K5NOSH polymers. 3D representations of the (B) heparin, (D) K5NOH, and (F) K5NOSH tetrasaccharides used to build the 31mer chains (with 31 monomers) bound to spike by employing the incremental sliding window docking method^1^. In the incremental sliding window docking method, docking begins with a 4-mer probe. The docking grid then shifts along the identified basic groove path, overlapping with the last monosaccharide of the docked 4-mer. This ensures that the last monosaccharide of one probe overlaps with the first monosaccharide of the next, preserving the glycosidic linkage properties between monosaccharides. With 10 iterations, this process results in a chain of 31 monosaccharides, with an extra monosaccharide at the end of the chain. Heparin, K5OSH, and K5NOSH are shown in stick representation colored by element with green, pink, and purple carbons, respectively.

### Supporting Tables

**Tab. S1.**
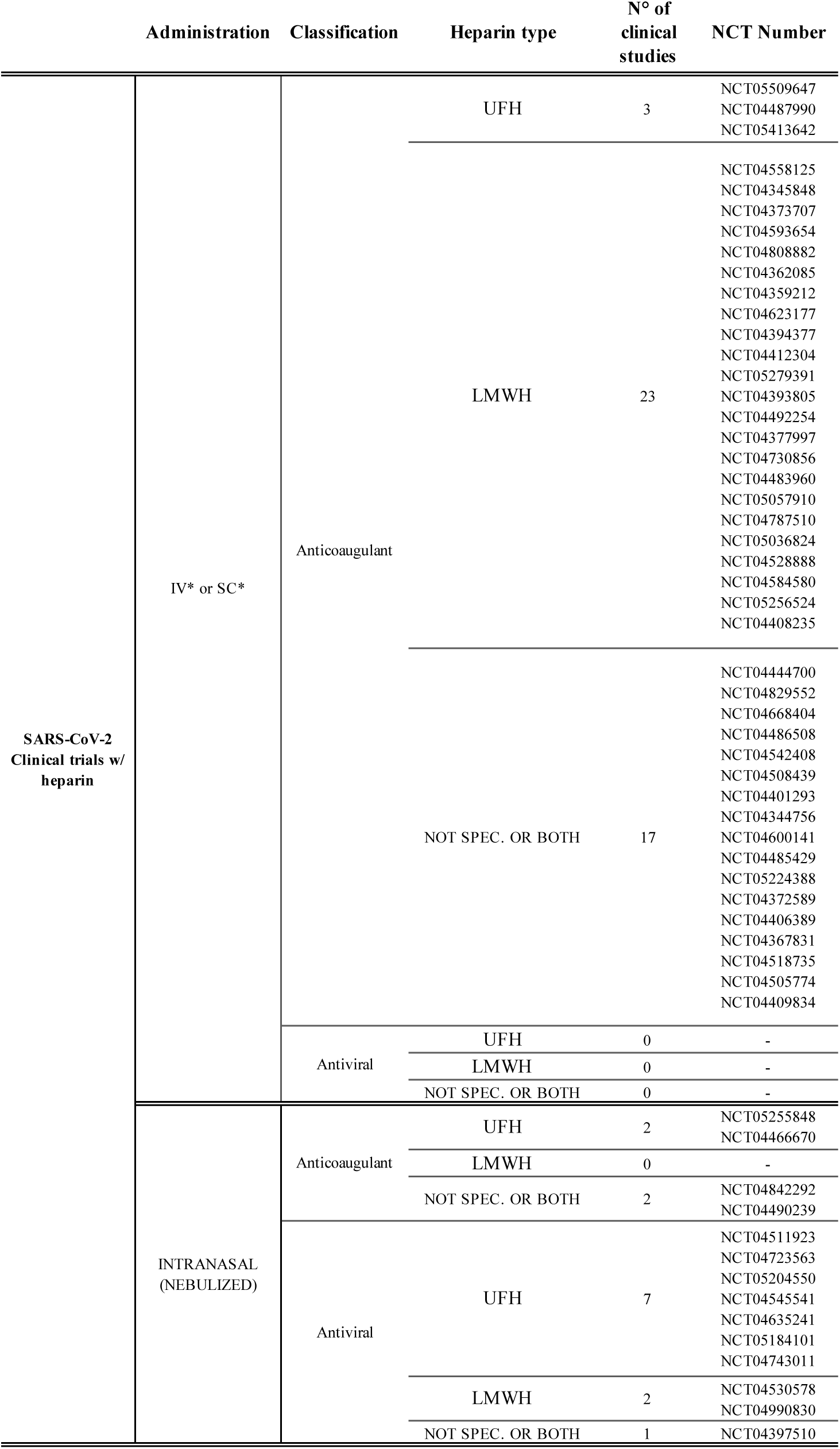
Clinical trials performed on Covid-19 patients using UFH or LMWH. The information was retrieved from clinicaltrials.gov with “Disease: Covid-19 Treatment: Heparin” and “Disease: Sars-Cov-2 Treatment: Heparin” as search items. The raw data summarized in the table are available on the Zenodo repository.

**Tab.S2.** ^13^C NMR spectrum analysis and SO_3_^-^/COO^-^ ratio analysis of the GAGs used in this work and retrieved from Casu et al^3^. na stands for not applicable because heparin does not have these monosaccharides in its structure.

| GAG | $\text{SO}_3^-/\text{COO}^-$<br>(ratio) | Glc-<br>$\text{NSO}_3^-$<br>(%) | Glc-<br>$6\text{SO}_3^-$<br>(%) | GlcA2$\text{SO}_3^-$ or<br>GlcA3$\text{SO}_3^-$<br>(%) | GlcA2,<br>3$\text{SO}_3^-$<br>(%) | unsulfated<br>GlcA<br>(%) | MW<br>(kDa) |
| --- | --- | --- | --- | --- | --- | --- | --- |
| Heparin | 2.14 | 78 | 77 | 59 | na | na | 13.6 |
| K5 | 0 | 0 | 0 | 0 | 0 | 100 | 30.0 |
| K5NS | 1.00 | 100 | 0 | 0 | 0 | 100 | 15.0 |
| K5OSL | 1.41 | 0 | 90 | <10 | 0 | >90 | 14.0 |
| K5NOSL | 1.70 | 100 | 90 | <10 | 0 | >90 | 13.0 |
| K5OSH | 3.77 | 0 | 100 | 0 | 100 | 0 | 11.0 |
| K5NOSH | 3.84 | 100 | 100 | 30 | 70 | 0 | 15.0 |

**Tab. S3.**
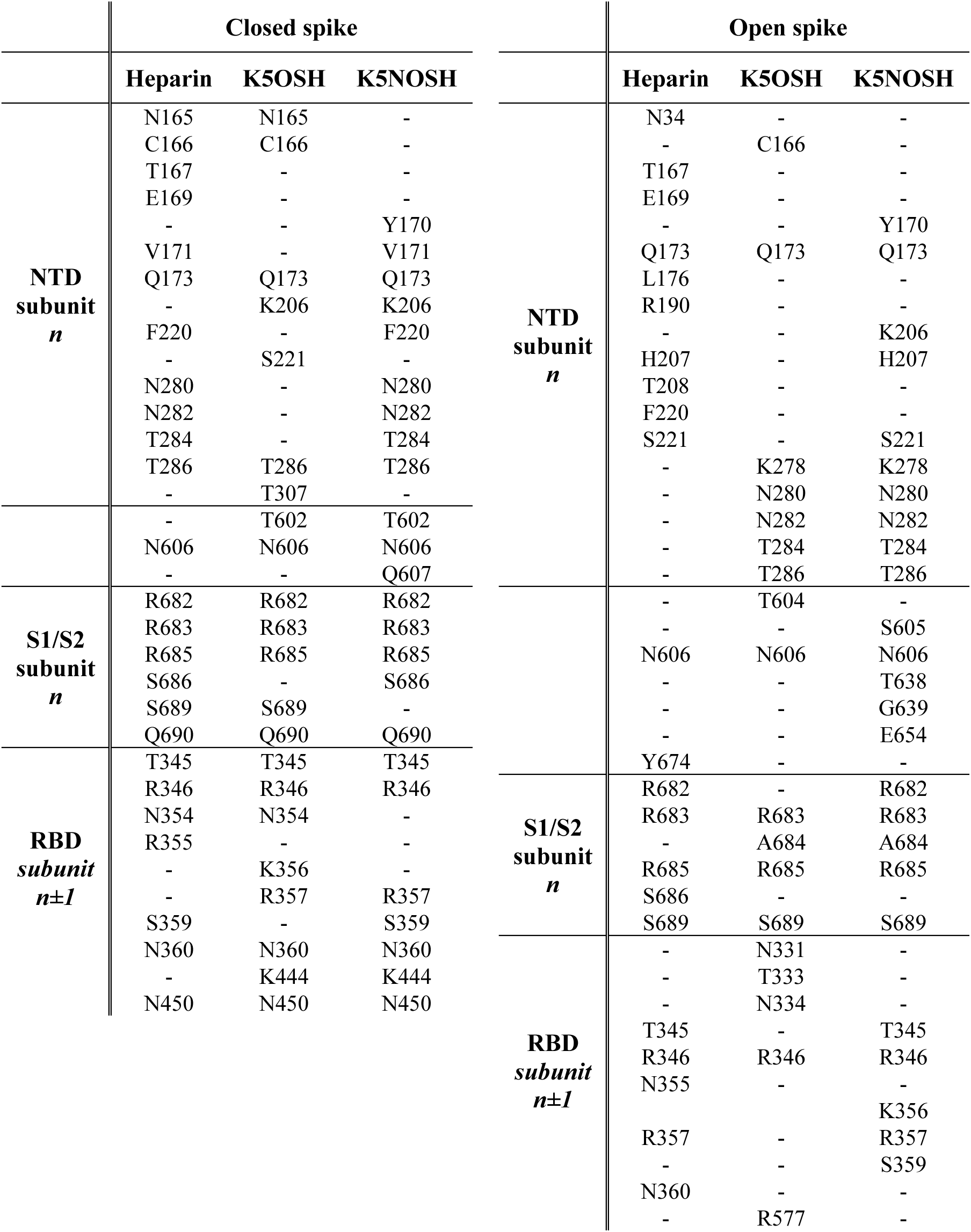
Residues of closed (left) and open (right) spike involved in hydrogen bonding interactions with heparin Paiardi et al.^2^, K5OSH and K5NOSH. H-bonds are reported when interacting in more than 75% of the trajectory and in at least 2 out of 4 replica simulations for each system.

**Tab. S4.** Analysis of hydrogen bonds (H-bonds) between closed spike glycoprotein and the K5 compounds for the four replica MD trajectories of the two simulated systems with K5OSH (top) or K5NOSH (bottom) compounds bound to closed spike. Each K5 chain interacts with two adjacent spike subunits as follows: K5OSH (chain 1) and K5NOSH (chain 1) interact with S1/S2 and RBD of S_A_ and S_B_, respectively; K5OSH (chain 2) and K5NOSH (chain 2) interact with S1/S2 and RBD of S_B_ and S_C_, respectively; K5OSH (chain 3) and K5NOSH (chain 3) interact with S1/S2 and RBD of S_C_ and S_A_, respectively. Residues involved in the interaction are listed separately as residues of the subunit *n* and the subunit *n±1*. The orange and pink boxes represent H-bonds with occupancy > 50% in a single trajectory and present in more than half the simulations with K5OSH or K5NOSH respectively.

| Closed spike +K5OSH |  |  |  |  |  |  |  |  |  |  |  |  |  |
| --- | --- | --- | --- | --- | --- | --- | --- | --- | --- | --- | --- | --- | --- |
|  | REPLICA 1 |  |  | REPLICA 2 |  |  | REPLICA 3 |  |  | REPLICA 4 |  |  |  |
| Subunit<br>$n$ | K5<br>OSH<br>(1) | K5<br>OSH<br>(2) | K5<br>OSH<br>(3) | K5<br>OSH<br>(1) | K5<br>OSH<br>(2) | K5<br>OSH<br>(3) | K5<br>OSH<br>(1) | K5<br>OSH<br>(2) | K5<br>OSH<br>(3) | K5<br>OSH<br>(1) | K5<br>OSH<br>(2) | K5<br>OSH<br>(3) | SUM |
| N164 | 0 | 1 | 1 | 0 | 1 | 0 | 0 | 0 | 0 | 1 | 0 | 0 | 2 |
| N165 | 1 | 1 | 1 | 1 | 1 | 1 | 0 | 1 | 1 | 1 | 1 | 1 | 11 |
| C166 | 1 | 0 | 1 | 0 | 1 | 1 | 0 | 1 | 0 | 0 | 0 | 1 | 6 |
| Y170 | 1 | 1 | 1 | 0 | 0 | 0 | 0 | 0 | 0 | 0 | 0 | 0 | 3 |
| V171 | 1 | 0 | 1 | 0 | 1 | 0 | 0 | 1 | 0 | 0 | 0 | 0 | 4 |
| Q173 | 0 | 1 | 1 | 0 | 1 | 0 | 0 | 1 | 1 | 0 | 1 | 0 | 6 |
| K206 | 1 | 0 | 0 | 1 | 0 | 0 | 0 | 0 | 1 | 1 | 1 | 1 | 6 |
| H207 | 0 | 0 | 0 | 0 | 0 | 1 | 1 | 0 | 0 | 0 | 1 | 0 | 3 |
| F220 | 0 | 0 | 0 | 0 | 1 | 0 | 0 | 0 | 0 | 1 | 0 | 0 | 2 |
| S221 | 0 | 1 | 0 | 0 | 1 | 1 | 0 | 1 | 1 | 0 | 1 | 0 | 6 |
| K278 | 0 | 0 | 0 | 0 | 0 | 0 | 1 | 0 | 0 | 0 | 0 | 1 | 2 |
| N280 | 0 | 0 | 0 | 0 | 1 | 0 | 0 | 0 | 1 | 1 | 0 | 1 | 4 |
| N282 | 0 | 0 | 0 | 0 | 1 | 1 | 0 | 0 | 0 | 0 | 0 | 1 | 3 |
| T284 | 0 | 0 | 0 | 0 | 0 | 0 | 0 | 1 | 0 | 1 | 0 | 0 | 2 |
| T286 | 1 | 1 | 0 | 1 | 0 | 0 | 1 | 1 | 0 | 0 | 0 | 1 | 6 |
| T307 | 0 | 1 | 0 | 1 | 1 | 0 | 1 | 1 | 0 | 0 | 0 | 1 | 6 |
| T602 | 0 | 1 | 0 | 1 | 1 | 0 | 1 | 1 | 0 | 0 | 0 | 1 | 6 |
| S605 | 0 | 0 | 0 | 0 | 1 | 0 | 0 | 1 | 0 | 0 | 1 | 0 | 3 |
| N606 | 1 | 1 | 0 | 0 | 1 | 0 | 1 | 1 | 1 | 0 | 1 | 1 | 8 |
| Q607 | 0 | 1 | 0 | 0 | 1 | 0 | 0 | 1 | 0 | 0 | 1 | 0 | 4 |
| P681 | 0 | 0 | 0 | 0 | 0 | 1 | 0 | 0 | 0 | 1 | 0 | 0 | 2 |
| R682 | 1 | 1 | 1 | 0 | 1 | 0 | 0 | 1 | 1 | 1 | 1 | 1 | 9 |
| R683 | 1 | 1 | 1 | 1 | 0 | 1 | 1 | 1 | 1 | 0 | 0 | 1 | 9 |
| R685 | 0 | 1 | 1 | 1 | 1 | 1 | 1 | 1 | 1 | 1 | 1 | 1 | 11 |
| S686 | 0 | 0 | 0 | 0 | 0 | 0 | 1 | 0 | 1 | 1 | 0 | 0 | 3 |
| S689 | 1 | 1 | 0 | 0 | 1 | 0 | 0 | 1 | 0 | 0 | 1 | 1 | 6 |
| Q690 | 1 | 1 | 0 | 0 | 1 | 0 | 0 | 1 | 0 | 0 | 1 | 1 | 6 |
| Subunit<br>$n\pm 1$ | K5<br>OSH<br>(1) | K5<br>OSH<br>(2) | K5<br>OSH<br>(3) | K5<br>OSH<br>(1) | K5<br>OSH<br>(2) | K5<br>OSH<br>(3) | K5<br>OSH<br>(1) | K5<br>OSH<br>(2) | K5<br>OSH<br>(3) | K5<br>OSH<br>(1) | K5<br>OSH<br>(2) | K5<br>OSH<br>(3) | SUM |
| T345 | 0 | 1 | 0 | 1 | 1 | 0 | 1 | 1 | 0 | 0 | 1 | 1 | 7 |
| R346 | 1 | 1 | 1 | 1 | 1 | 1 | 1 | 1 | 1 | 1 | 1 | 1 | 12 |
| N354 | 1 | 0 | 0 | 1 | 1 | 0 | 1 | 1 | 0 | 1 | 0 | 0 | 6 |
| K356 | 0 | 1 | 1 | 1 | 1 | 0 | 1 | 1 | 0 | 1 | 0 | 1 | 8 |
| R357 | 0 | 1 | 1 | 1 | 1 | 1 | 1 | 1 | 1 | 1 | 1 | 1 | 11 |
| S359 | 0 | 1 | 0 | 0 | 0 | 1 | 0 | 0 | 1 | 1 | 0 | 0 | 4 |
| N360 | 1 | 1 | 1 | 0 | 1 | 1 | 0 | 1 | 1 | 1 | 1 | 1 | 10 |
| N394 | 0 | 1 | 0 | 0 | 0 | 0 | 1 | 1 | 1 | 0 | 0 | 0 | 4 |
| K444 | 1 | 1 | 0 | 1 | 1 | 1 | 1 | 0 | 0 | 0 | 1 | 1 | 8 |
| V445 | 1 | 1 | 0 | 0 | 0 | 0 | 0 | 0 | 1 | 0 | 0 | 0 | 3 |
| G447 | 1 | 1 | 0 | 1 | 1 | 0 | 0 | 0 | 0 | 0 | 0 | 0 | 4 |
| N448 | 0 | 0 | 0 | 0 | 0 | 0 | 0 | 0 | 1 | 0 | 0 | 1 | 2 |
| Y449 | 1 | 1 | 0 | 1 | 1 | 1 | 0 | 0 | 0 | 0 | 0 | 0 | 5 |
| N450 | 1 | 0 | 0 | 0 | 1 | 1 | 1 | 0 | 1 | 0 | 0 | 1 | 6 |

| Closed spike +K5NOSH |  |  |  |  |  |  |  |  |  |  |  |  |  |
| --- | --- | --- | --- | --- | --- | --- | --- | --- | --- | --- | --- | --- | --- |
|  | REPLICA 1 |  |  | REPLICA 2 |  |  | REPLICA 3 |  |  | REPLICA 4 |  |  |  |
| Subunit<br>*n* | K5<br>NOSH<br>(1) | K5<br>NOSH<br>(2) | K5<br>NOSH<br>(3) | K5<br>NOSH<br>(1) | K5<br>NOSH<br>(2) | K5<br>NOSH<br>(3) | K5<br>NOSH<br>(1) | K5<br>NOSH<br>(2) | K5<br>NOSH<br>(3) | K5<br>NOSH<br>(1) | K5<br>NOSH<br>(2) | K5<br>NOSH<br>(3) | SUM |
| N164 | 1 | 0 | 0 | 0 | 0 | 0 | 0 | 0 | 0 | 0 | 1 | 1 | 3 |
| N165 | 1 | 0 | 0 | 0 | 0 | 0 | 0 | 0 | 0 | 0 | 0 | 1 | 2 |
| E169 | 0 | 0 | 0 | 1 | 1 | 1 | 0 | 0 | 0 | 0 | 1 | 0 | 4 |
| Y170 | 1 | 1 | 0 | 1 | 1 | 0 | 1 | 1 | 0 | 1 | 1 | 1 | 9 |
| V171 | 0 | 1 | 0 | 1 | 0 | 0 | 0 | 1 | 1 | 1 | 0 | 1 | 6 |
| S172 | 0 | 1 | 0 | 1 | 0 | 0 | 0 | 0 | 1 | 1 | 0 | 0 | 4 |
| Q173 | 1 | 1 | 1 | 1 | 1 | 1 | 1 | 1 | 1 | 0 | 0 | 1 | 10 |
| Q183 | 1 | 0 | 0 | 0 | 0 | 0 | 0 | 0 | 0 | 0 | 0 | 1 | 2 |
| K206 | 0 | 1 | 1 | 0 | 1 | 0 | 0 | 1 | 1 | 0 | 0 | 1 | 6 |
| H207 | 0 | 0 | 1 | 0 | 0 | 0 | 0 | 1 | 1 | 0 | 0 | 1 | 4 |
| Q218 | 1 | 0 | 0 | 0 | 1 | 1 | 0 | 0 | 0 | 0 | 0 | 0 | 3 |
| G219 | 0 | 0 | 0 | 0 | 1 | 1 | 0 | 0 | 0 | 0 | 1 | 0 | 3 |
| F220 | 1 | 1 | 0 | 1 | 1 | 1 | 1 | 1 | 0 | 0 | 1 | 0 | 8 |
| K278 | 0 | 0 | 1 | 0 | 0 | 0 | 0 | 1 | 0 | 0 | 0 | 0 | 2 |
| N280 | 0 | 1 | 1 | 1 | 0 | 1 | 1 | 1 | 1 | 0 | 0 | 1 | 8 |
| N282 | 1 | 1 | 1 | 1 | 0 | 1 | 1 | 0 | 1 | 0 | 0 | 1 | 8 |
| T284 | 0 | 1 | 1 | 1 | 1 | 1 | 0 | 1 | 0 | 1 | 1 | 0 | 8 |
| T286 | 0 | 1 | 0 | 0 | 1 | 1 | 1 | 1 | 1 | 0 | 0 | 1 | 7 |
| S605 | 0 | 1 | 0 | 0 | 0 | 0 | 1 | 0 | 0 | 1 | 0 | 0 | 3 |
| N606 | 0 | 1 | 0 | 1 | 0 | 0 | 1 | 1 | 1 | 1 | 0 | 0 | 6 |
| Q607 | 1 | 1 | 0 | 0 | 1 | 0 | 0 | 1 | 0 | 1 | 1 | 0 | 6 |
| S637 | 0 | 1 | 0 | 0 | 1 | 0 | 0 | 0 | 0 | 0 | 0 | 0 | 2 |
| T638 | 0 | 1 | 0 | 0 | 1 | 0 | 0 | 1 | 0 | 0 | 0 | 0 | 3 |
| S640 | 0 | 1 | 0 | 0 | 0 | 0 | 0 | 0 | 0 | 1 | 0 | 0 | 2 |
| E654 | 0 | 1 | 0 | 1 | 0 | 0 | 0 | 0 | 0 | 1 | 0 | 0 | 3 |
| S680 | 1 | 0 | 0 | 0 | 0 | 1 | 0 | 0 | 0 | 0 | 0 | 0 | 2 |
| P681 | 0 | 0 | 0 | 0 | 0 | 1 | 0 | 0 | 0 | 1 | 0 | 0 | 2 |
| R682 | 1 | 1 | 0 | 1 | 0 | 1 | 1 | 1 | 0 | 1 | 1 | 1 | 9 |
| R683 | 1 | 0 | 0 | 1 | 1 | 1 | 1 | 1 | 1 | 1 | 0 | 0 | 8 |
| R685 | 1 | 1 | 0 | 1 | 1 | 0 | 1 | 0 | 1 | 0 | 1 | 1 | 8 |
| S686 | 1 | 0 | 0 | 1 | 1 | 0 | 1 | 0 | 0 | 1 | 0 | 1 | 6 |
| S689 | 1 | 0 | 0 | 0 | 1 | 1 | 0 | 0 | 0 | 0 | 0 | 0 | 3 |
| Q690 | 1 | 1 | 0 | 1 | 0 | 1 | 1 | 1 | 0 | 0 | 0 | 0 | 6 |
| S691 | 0 | 1 | 0 | 0 | 1 | 1 | 0 | 0 | 0 | 1 | 0 | 0 | 4 |
| Subunit<br>*n±1* | K5<br>NOSH<br>(1) | K5<br>NOSH<br>(2) | K5<br>NOSH<br>(3) | K5<br>NOSH<br>(1) | K5<br>NOSH<br>(2) | K5<br>NOSH<br>(3) | K5<br>NOSH<br>(1) | K5<br>NOSH<br>(2) | K5<br>NOSH<br>(3) | K5<br>NOSH<br>(1) | K5<br>NOSH<br>(2) | K5<br>NOSH<br>(3) | SUM |
| N334 | 0 | 0 | 0 | 1 | 0 | 0 | 1 | 0 | 0 | 0 | 0 | 0 | 2 |
| T345 | 0 | 1 | 0 | 1 | 0 | 1 | 1 | 0 | 0 | 0 | 1 | 1 | 6 |
| R346 | 0 | 1 | 1 | 1 | 1 | 1 | 1 | 1 | 1 | 1 | 1 | 1 | 11 |
| N354 | 0 | 0 | 0 | 0 | 1 | 0 | 0 | 0 | 1 | 0 | 0 | 0 | 2 |
| K356 | 0 | 1 | 0 | 0 | 0 | 0 | 0 | 1 | 0 | 0 | 1 | 1 | 4 |
| R357 | 1 | 1 | 1 | 1 | 1 | 1 | 1 | 1 | 1 | 1 | 1 | 0 | 11 |
| S359 | 0 | 1 | 1 | 0 | 1 | 1 | 0 | 0 | 1 | 1 | 0 | 1 | 7 |
| N360 | 1 | 1 | 1 | 0 | 1 | 1 | 1 | 1 | 1 | 1 | 0 | 1 | 10 |
| K444 | 0 | 1 | 0 | 1 | 1 | 1 | 0 | 1 | 0 | 1 | 0 | 0 | 6 |
| K449 | 0 | 0 | 0 | 0 | 0 | 0 | 0 | 1 | 0 | 1 | 0 | 0 | 2 |
| N450 | 0 | 1 | 0 | 1 | 1 | 1 | 1 | 0 | 0 | 1 | 1 | 0 | 7 |

**Tab. S5.** Analysis of hydrogen bonds (H-bonds) between open spike glycoprotein and the K5 compounds for the four replica MD trajectories of the two simulated systems with open spike and K5OSH or K5NOSH compounds bound. Each K5 chain interacts with two adjacent spike subunits as follows: K5OSH (chain 1) and K5NOSH (chain 1) interact with S1/S2 and RBD of S_A_ and S_B_, respectively; K5OSH (chain 2) and K5NOSH (chain 2) interact with S1/S2 and RBD of S_B_ and S_C_, respectively; K5OSH (chain 3) and K5NOSH (chain 3) interacts with S1/S2 and RBD of S_C_ and S_A_, respectively. Residues involved in the interaction are listed separately as residues of the subunit *n* and the subunit n±1. The orange and pink boxes represent the H-bonds with occupancy > 50% in a single trajectory and stable in more than half the simulations on the subunits with K5OSH or K5NOSH respectively, considering the single chain (chain 2) on the open subunit (SUM open) or the two K5 chains (chain 1 and chain 3) on the closed subunits (SUM closed).

| Open spike +K5OSH |  |  |  |  |  |  |  |  |  |  |  |  |  |  |
| --- | --- | --- | --- | --- | --- | --- | --- | --- | --- | --- | --- | --- | --- | --- |
|  | REPLICA 1 |  |  | REPLICA 2 |  |  | REPLICA 3 |  |  | REPLICA 4 |  |  |  |  |
| Subunit $n$ | K5 OSH (1) | K5 OSH (2) | K5 OSH (3) | K5 OSH (1) | K5 OSH (2) | K5 OSH (3) | K5 OSH (1) | K5 OSH (2) | K5 OSH (3) | K5 OSH (1) | K5 OSH (2) | K5 OSH (3) | SUM open | SUM closed |
| N164 | 0 | 0 | 0 | 0 | 0 | 1 | 1 | 0 | 0 | 0 | 0 | 0 | 0 | 2 |
| N165 | 1 | 1 | 0 | 0 | 0 | 1 | 1 | 0 | 1 | 1 | 0 | 0 | 1 | 5 |
| C166 | 0 | 0 | 0 | 0 | 1 | 0 | 0 | 0 | 0 | 0 | 1 | 0 | 2 | 0 |
| T167 | 1 | 0 | 0 | 1 | 0 | 0 | 0 | 0 | 1 | 0 | 0 | 0 | 0 | 3 |
| F168 | 1 | 0 | 1 | 0 | 0 | 0 | 0 | 0 | 1 | 0 | 0 | 0 | 0 | 3 |
| Q173 | 0 | 1 | 1 | 0 | 0 | 0 | 1 | 1 | 1 | 0 | 1 | 0 | 3 | 3 |
| K206 | 1 | 0 | 1 | 0 | 0 | 1 | 0 | 0 | 1 | 0 | 0 | 0 | 0 | 4 |
| H207 | 0 | 0 | 1 | 0 | 0 | 1 | 0 | 0 | 1 | 0 | 0 | 0 | 0 | 3 |
| T208 | 0 | 0 | 1 | 0 | 0 | 1 | 1 | 0 | 1 | 0 | 0 | 1 | 0 | 5 |
| R214 | 0 | 0 | 0 | 0 | 0 | 1 | 0 | 0 | 0 | 1 | 0 | 1 | 0 | 3 |
| S221 | 0 | 0 | 0 | 0 | 1 | 0 | 1 | 0 | 0 | 0 | 0 | 1 | 1 | 2 |
| K278 | 0 | 1 | 1 | 0 | 0 | 0 | 0 | 1 | 0 | 0 | 0 | 0 | 2 | 1 |
| N280 | 0 | 0 | 1 | 1 | 0 | 1 | 0 | 1 | 1 | 0 | 1 | 1 | 2 | 5 |
| N282 | 0 | 0 | 0 | 1 | 0 | 0 | 0 | 1 | 0 | 1 | 1 | 0 | 2 | 2 |
| T284 | 0 | 0 | 0 | 0 | 0 | 0 | 0 | 1 | 0 | 0 | 1 | 0 | 2 | 0 |
| T286 | 0 | 1 | 0 | 0 | 0 | 1 | 1 | 1 | 0 | 0 | 1 | 0 | 3 | 2 |
| T604 | 0 | 1 | 0 | 0 | 1 | 0 | 0 | 1 | 0 | 0 | 1 | 0 | 4 | 0 |
| N606 | 0 | 0 | 1 | 1 | 1 | 1 | 0 | 1 | 1 | 0 | 0 | 0 | 2 | 4 |
| S680 | 0 | 0 | 1 | 0 | 0 | 0 | 0 | 1 | 0 | 1 | 0 | 0 | 1 | 2 |
| R682 | 1 | 0 | 1 | 1 | 0 | 1 | 1 | 0 | 1 | 1 | 1 | 1 | 1 | 8 |
| R683 | 1 | 1 | 1 | 1 | 1 | 1 | 1 | 1 | 1 | 1 | 1 | 1 | 4 | 8 |
| A684 | 1 | 1 | 0 | 0 | 1 | 0 | 0 | 1 | 1 | 0 | 0 | 0 | 3 | 2 |
| R685 | 0 | 1 | 1 | 1 | 1 | 1 | 0 | 1 | 1 | 0 | 1 | 1 | 4 | 5 |
| S686 | 1 | 0 | 1 | 0 | 1 | 0 | 0 | 0 | 0 | 0 | 0 | 1 | 1 | 3 |
| S689 | 0 | 0 | 0 | 0 | 0 | 0 | 0 | 1 | 0 | 0 | 1 | 0 | 2 | 0 |
| Q690 | 0 | 0 | 1 | 0 | 1 | 1 | 0 | 0 | 1 | 0 | 0 | 0 | 1 | 3 |
| Subunit $n\pm 1$ | K5 OSH1 | K5 OSH2 | K5 OSH3 | K5 OSH1 | K5 OSH2 | K5 OSH3 | K5 OSH1 | K5 OSH2 | K5 OSH3 | K5 OSH1 | K5 OSH2 | K5 OSH3 | SUM open | SUM closed |
| N331 | 0 | 0 | 0 | 0 | 0 | 0 | 0 | 1 | 0 | 0 | 1 | 0 | 2 | 0 |
| T333 | 0 | 0 | 0 | 0 | 0 | 1 | 0 | 1 | 0 | 0 | 1 | 0 | 2 | 1 |
| N334 | 0 | 1 | 0 | 0 | 1 | 1 | 1 | 1 | 0 | 0 | 1 | 0 | 4 | 2 |
| R346 | 1 | 1 | 1 | 1 | 1 | 1 | 1 | 0 | 1 | 1 | 1 | 1 | 3 | 8 |
| N354 | 0 | 0 | 1 | 0 | 0 | 0 | 0 | 0 | 0 | 0 | 0 | 1 | 0 | 2 |
| K356 | 0 | 0 | 0 | 0 | 0 | 1 | 0 | 0 | 1 | 0 | 1 | 0 | 1 | 2 |
| R357 | 1 | 0 | 1 | 0 | 0 | 1 | 0 | 0 | 1 | 1 | 0 | 1 | 0 | 6 |
| S359 | 1 | 0 | 1 | 1 | 0 | 0 | 0 | 0 | 0 | 1 | 0 | 0 | 0 | 4 |
| N360 | 0 | 0 | 1 | 1 | 0 | 1 | 0 | 0 | 0 | 1 | 0 | 1 | 0 | 5 |
| K444 | 1 | 0 | 0 | 0 | 0 | 0 | 1 | 0 | 0 | 0 | 1 | 0 | 1 | 2 |
| K558 | 0 | 1 | 0 | 1 | 0 | 0 | 0 | 0 | 0 | 0 | 0 | 1 | 1 | 2 |
| R577 | 0 | 1 | 0 | 0 | 1 | 0 | 0 | 1 | 0 | 0 | 0 | 0 | 3 | 0 |

| Open spike +K5NOSH |  |  |  |  |  |  |  |  |  |  |  |  |  |  |
| --- | --- | --- | --- | --- | --- | --- | --- | --- | --- | --- | --- | --- | --- | --- |
|  | REPLICA 1 |  |  | REPLICA 2 |  |  | REPLICA 3 |  |  | REPLICA 4 |  |  |  |  |
| Subunit<br>*n* | K5<br>NOSH<br>(1) | K5<br>NOSH<br>(2) | K5<br>NOSH<br>(3) | K5<br>NOSH<br>(1) | K5<br>NOSH<br>(2) | K5<br>NOSH<br>(3) | K5<br>NOSH<br>(1) | K5<br>NOSH<br>(2) | K5<br>NOSH<br>(3) | K5<br>NOSH<br>(1) | K5<br>NOSH<br>(2) | K5<br>NOSH<br>(3) | SUM<br>open | SUM<br>closed |
| N125 | 0 | 0 | 0 | 0 | 0 | 0 | 0 | 0 | 1 | 0 | 0 | 1 | 0 | 2 |
| C166 | 0 | 0 | 0 | 0 | 0 | 0 | 1 | 0 | 1 | 0 | 0 | 0 | 0 | 2 |
| Y170 | 0 | 1 | 0 | 0 | 1 | 0 | 0 | 0 | 0 | 0 | 0 | 0 | 2 | 0 |
| Q173 | 1 | 0 | 1 | 0 | 1 | 1 | 1 | 1 | 1 | 1 | 0 | 1 | 2 | 7 |
| K206 | 0 | 1 | 0 | 1 | 1 | 0 | 0 | 1 | 0 | 1 | 1 | 0 | 4 | 2 |
| H207 | 0 | 0 | 1 | 0 | 1 | 0 | 0 | 1 | 1 | 0 | 1 | 1 | 3 | 3 |
| F220 | 0 | 0 | 1 | 0 | 0 | 1 | 1 | 0 | 1 | 0 | 0 | 0 | 0 | 4 |
| S221 | 0 | 1 | 0 | 1 | 1 | 1 | 0 | 0 | 1 | 1 | 0 | 0 | 2 | 4 |
| K278 | 0 | 0 | 0 | 0 | 0 | 0 | 0 | 1 | 0 | 0 | 1 | 1 | 2 | 1 |
| N280 | 0 | 1 | 0 | 0 | 1 | 0 | 1 | 1 | 0 | 1 | 1 | 1 | 4 | 3 |
| N282 | 0 | 1 | 1 | 1 | 1 | 0 | 0 | 0 | 0 | 1 | 0 | 1 | 2 | 4 |
| T284 | 0 | 1 | 0 | 1 | 1 | 0 | 0 | 1 | 1 | 1 | 0 | 1 | 3 | 4 |
| T286 | 0 | 1 | 0 | 0 | 1 | 0 | 1 | 1 | 1 | 1 | 0 | 0 | 3 | 3 |
| S605 | 0 | 0 | 0 | 0 | 1 | 0 | 1 | 1 | 1 | 1 | 0 | 0 | 2 | 3 |
| N606 | 0 | 1 | 1 | 0 | 1 | 1 | 0 | 1 | 1 | 0 | 0 | 0 | 3 | 3 |
| Q607 | 0 | 0 | 0 | 0 | 0 | 1 | 0 | 0 | 1 | 1 | 1 | 1 | 1 | 4 |
| T638 | 0 | 0 | 0 | 0 | 0 | 0 | 0 | 1 | 0 | 0 | 1 | 0 | 2 | 0 |
| G639 | 0 | 0 | 0 | 0 | 0 | 0 | 0 | 1 | 0 | 0 | 1 | 0 | 2 | 0 |
| E654 | 0 | 0 | 0 | 0 | 0 | 0 | 0 | 1 | 0 | 0 | 1 | 0 | 2 | 0 |
| R682 | 0 | 1 | 1 | 1 | 1 | 1 | 1 | 0 | 1 | 1 | 0 | 0 | 2 | 6 |
| R683 | 1 | 1 | 1 | 1 | 1 | 1 | 1 | 1 | 1 | 1 | 1 | 1 | 4 | 8 |
| A684 | 1 | 0 | 1 | 1 | 1 | 1 | 0 | 0 | 0 | 0 | 1 | 0 | 2 | 4 |
| R685 | 1 | 1 | 1 | 0 | 1 | 1 | 1 | 1 | 1 | 1 | 1 | 1 | 4 | 7 |
| S686 | 1 | 0 | 1 | 0 | 0 | 0 | 0 | 0 | 0 | 1 | 0 | 1 | 0 | 4 |
| V687 | 1 | 0 | 0 | 1 | 0 | 0 | 0 | 0 | 0 | 1 | 0 | 0 | 0 | 3 |
| A688 | 1 | 0 | 0 | 1 | 0 | 0 | 1 | 0 | 0 | 1 | 0 | 0 | 0 | 4 |
| S689 | 0 | 0 | 0 | 0 | 0 | 0 | 1 | 1 | 1 | 0 | 1 | 0 | 2 | 2 |
| Q690 | 1 | 1 | 0 | 1 | 1 | 1 | 1 | 0 | 1 | 1 | 1 | 0 | 3 | 6 |
| S691 | 1 | 0 | 0 | 0 | 0 | 1 | 1 | 0 | 0 | 0 | 0 | 0 | 0 | 3 |
| Subunit<br>*n*±1 | K5<br>NOSH<br>(1) | K5<br>NOSH<br>(2) | K5<br>NOSH<br>(3) | K5<br>NOSH<br>(1) | K5<br>NOSH<br>(2) | K5<br>NOSH<br>(3) | K5<br>NOSH<br>(1) | K5<br>NOSH<br>(2) | K5<br>NOSH<br>(3) | K5<br>NOSH<br>(1) | K5<br>NOSH<br>(2) | K5<br>NOSH<br>(3) | SUM<br>open | SUM<br>closed |
| N334 | 1 | 0 | 1 | 0 | 0 | 0 | 0 | 0 | 0 | 0 | 0 | 1 | 0 | 3 |
| T345 | 1 | 1 | 0 | 0 | 1 | 0 | 1 | 1 | 0 | 1 | 1 | 1 | 4 | 4 |
| R346 | 1 | 1 | 0 | 1 | 1 | 1 | 1 | 1 | 1 | 1 | 1 | 1 | 4 | 7 |
| F347 | 0 | 0 | 0 | 0 | 0 | 0 | 1 | 0 | 0 | 0 | 0 | 1 | 0 | 2 |
| K356 | 1 | 1 | 1 | 0 | 1 | 0 | 0 | 1 | 1 | 1 | 0 | 1 | 3 | 5 |
| R357 | 1 | 1 | 1 | 1 | 1 | 1 | 1 | 1 | 1 | 1 | 0 | 1 | 3 | 8 |
| S359 | 1 | 0 | 1 | 1 | 1 | 1 | 1 | 1 | 1 | 0 | 0 | 0 | 2 | 6 |
| N360 | 1 | 0 | 1 | 1 | 0 | 1 | 1 | 1 | 1 | 1 | 0 | 1 | 1 | 8 |
| K444 | 1 | 0 | 0 | 0 | 0 | 0 | 1 | 0 | 1 | 0 | 1 | 0 | 1 | 3 |
| N448 | 0 | 0 | 0 | 0 | 0 | 0 | 1 | 0 | 1 | 0 | 1 | 0 | 1 | 3 |
| N450 | 0 | 0 | 0 | 0 | 0 | 1 | 1 | 0 | 1 | 0 | 0 | 1 | 0 | 4 |

